# The complete chloroplast genome sequence of American elm (*Ulmus americana*) and comparative genomics of related species

**DOI:** 10.1101/2020.01.27.920181

**Authors:** Aziz Ebrahimi, Jennifer D. Antonides, Cornelia C. Pinchot, James M. Slavicek, Charles E. Flower, Keith E. Woeste

## Abstract

American elm, *Ulmus americana* L., was cultivated widely in USA and Canada as a landscape tree, but the genome of this important species is poorly characterized. For the first time, we describe the sequencing and assembly of the chloroplast genomes of two American elm genotypes (RV16 and Am57845). The complete chloroplast genome of *U. americana* ranged from 158,935-158,993 bp. The genome contains 127 genes, including 85 protein-coding genes, 34 tRNA genes and 8 rRNA genes. Between the two American elm chloroplasts we sequenced, we identified 240 sequence variants (SNPs and indels). To evaluate the phylogeny of American elm, we compared the chloroplast genomes of two American elms along with seven Asian elm species and twelve other chloroplast genomes available through the NCBI database. As expected, *Ulmus* was closely related to *Morus* and *Cannabis*, as all three genera are assigned to the Urticales. Comparison of American elm with Asian elms revealed that *trn*H was absent from the chloroplast of American elm but not most Asian elms; conversely, *pet*B, *pet*D, *psb*L, *trn*K, and *rps*16 are present in the American elm but absent from all Asian elms. The complete chloroplast genome of *U. americana* will provide useful genetic resources for characterizing the genetic diversity of *U. americana* and potentially help to conserve natural populations of American elm.

## INTRODUCTION

The American elm, *Ulmus americana* L. occurs as diploid (2n = 2x = 28), a tetraploid (2n = 4x = 56), and even as a triploid in its native range in the eastern United States and Canada (Bey, 1990;Whittemore and Xia, 2017). Of the six species of elm native to North America, *U. americana* has the largest historical range (NRCS, 2017), and it has been widely planted as an ornamental tree in many parts of the United States outside its native range. Exotic insects or pathogens have affected many American elm trees (Whittemore and Olsen, 2011). The most consequential is Dutch Elm Disease (DED), which, as described by Schwarz (1922) and Karnosky (1979), arose in the 20th century in USA and resulted in the death of a majority of mature elms across much of the northern hemisphere (Brasier and Buck, 2001). The remarkable expansion of DED led some to forecast the complete extirpation of elms in North America or, at best, its persistence as a scrub species (Brasier, 1983). Although the number of elms lost to DED since the epidemic began is uncertain, it is likely many millions of trees based on their former abundance (Bey, 1990). The changes in age structure associated with mortality of mature elms and the reduction in population sizes associated with DED could be expected to modify how genetic variation in elms is distributed over the landscape (Bouzat, 2010). Understanding these changes in the population genetics of elms may improve our ability to control the diseases of elm and to breed elms for the wide variety of uses for which they were once prized (Pinchot et al., 2017).

Chloroplast genomes can be used to understand evolution, genetic diversity and genetic structure because of their highly conserved structure, low rate of mutation, and uniparental inheritance (Setohigashi et al., 2011). Sequence data from chloroplast genomes have been used in phylogenetic analysis (Ruhfel et al., 2014), biotechnological applications (Sabir et al., 2014), species identification (Kane et al., 2012), identifying Single-nucleotide polymorphisms (SNPs), and studies of inter- and intraspecific variability (Alexander and Woeste, 2014). To date, seven complete chloroplast genomes in the genus *Ulmus* from Asia have been reported. These include: *U. pumila*., *U. laciniata*, *U. davidiana*, *U. macrocarpa*, *U. davidiana* var. *japonica* (Zuo et al., 2017), *U. gaussenii* (Zhang et al., 2019a), and *U. chenmoui* (Zhang et al., 2019b). However, the genomes of North American elms, including the most important North American species, *U. americana*, are not yet published. Previous phylogenetic analyses of *Ulmus* were based on SSR markers (Brunet and Guries, 2016) or variation level of ploidy in elm species (Whittemore and Olsen, 2011). Here, we report the genome sequence of two American elms based on paired-end whole genome sequencing. The main aims of the present study were (1) assemble the whole chloroplast genome of American elm, (2) evaluate the phylogenetic position of American elm relative to Asian elms, and (3) Identify genic and intergenic regions of polymorphism within the American elm chloroplast genome.

## MATERIALS AND METHODS

### Plant samples and sequencing

Fresh leaf samples of two genotypes (RV16 and Am. 57845) of *U. americana* were collected at the U.S.D.A. Forest Service, Northern Research Station work unit in Delaware, Ohio, USA. Genomic DNA was extracted using the methods of Zhao and Woeste (2011). Paired-end libraries were prepared by the Purdue Genomics Core Facility using the Truseq DNA PCR-Free Sample Preparation Kit (Illumina). Whole-genome sequencing was performed using Illumina short-read technology (2×100bp) on the HiSeq2500 sequencing platform in two single-lane reactions. Raw reads were processed using Trimmomatic (Bolger et al., 2014) to remove Illumina adapters and for quality control: bases of less than Phred-20 were clipped from the 5’ and 3’ ends, and subsequently reads below a minimum length of 30bp were discarded.

### Chloroplast genome assembly and annotation

The chloroplast genomes were assembled with BWA (Burrows Wheeler-aligner), Picard-tools and GATK (Genome Analysis Toolkit) software (Langmead et al., 2009;McKenna et al., 2010;Wysoker et al., 2019). The *U. davidiana* (NC_032718.1) genome (Zuo et al., 2017) was used as a preliminary reference to guide assembly of the RV16 genotype, and subsequently the RV16 assembly was used to fill gaps of the Am. 57845 genotype.

The RV16 assembly was annotated in stages; first by predicting genes using Mfannot (Beck and Lang, 2010), an integration of tools including BLAST (Altschul et al., 1990), Exonerate (Slater and Birney, 2005), RNAweasel (Lang et al., 2007) and collections of reference proteins, RNA models, and group I and II splice junction profiles. These predictions were used as references for annotation within the program GeSeq (Tillich et al., 2017), along with six publicly available chloroplast genomes in genus Ulmus: *U. chenmoui* (NC_037758.1), *U. davidiana* (NC_032718.1), *U. gaussenii* (NC_037840.1), *U. laciniata* (NC_032719.1), *U. macrocarpia* (NC_032720.1), and *U. pumila* (NC_032721.1), and GeSeq’s manually curated chloroplast rRNA and CDS reference sequences. The BLAT (Kent, 2002) search identity was set at 55% for protein and 85% for rRNA, tRNA, and DNA. The tRNA annotator ARAGORN (v. 1.2.38) was additionally used (Laslett and Canback, 2004). The annotations were validated in Sequin (v. 16.0), in which errors and warnings were flagged and resolved (NCBI, 2016). Subsequently the annotations were manually inspected to resolve discrepancies. As the strand directionality of some of the tRNAs and rRNAs in the *Ulmus* (Asian elm) chloroplast references were determined to be incorrect during annotation, the chloroplast genome of *Vitus vinifera* (NC_007957.1) was used as a reference to correct these issues. Subsequently the Am. 57845 assembly was annotated in a similar manner using the RV16 genome as an additional reference. The annotated American elm chloroplast genomes were deposited in the NCBI database (Genotype RV16 as MN043961 and Am. 57845 as MH324448).

### Analysis of U. americana chloroplast genomes

For both American elm genotypes a circular chloroplast genome map was drawn with OGDRAW (v. 1.3.1) (Greiner et al., 2019). A global pairwise sequence alignment of the two chloroplast genome sequences was produced using EMBOSS Stretcher (v. 6.6.0) with default parameters (Madeira et al., 2019). The alignment was visualized with the Sequence Manipulation Suite (Stothard, 2000).

### Comparison of American elm and Asian elms

The RV16 genotype was used as the representative *Ulmus americana* chloroplast genome, and compared to the chloroplasts of the seven Asian elm species publicly available (*U. pumila*., *U. laciniata*, *U. davidiana*, *U. macrocarpa*, *U. davidiana* var. *japonica*, *U. gaussenii*, and *U. chenmoui*). A global multiple sequence alignment of the eight species’ chloroplast genomes was produced using the LAGAN algorithm with translated anchoring (Brudno et al., 2003). The alignment, annotated according to the RV16 reference, was visualized with the mVISTA server (Frazer et al., 2004). The sizes and junction sites of the two inverted repeat (IR) regions, the short single-copy (SSC) region, and long single-copy (LSC) region were compared and visually represented with IRscope (Amiryousefi et al., 2018).

### Variant identification within the American elm chloroplast

To identify variants (i.e., SNPs and indels) within the *U. americana* chloroplast, chloroplast-specific reads were extracted from each genotype’s whole-genome paired-end reads using BBsplit in BBMap (v. 37.93) (Bushnell, 2014). These chloroplast reads from each genotype were separately mapped to the RV16 genotype’s chloroplast assembly using BWA (v. 0.7.15) (Langmead et al., 2009). Picardtools (v. 2.18.2) was used to sort and validate .sam and .bam files and to mark duplicates (Wysoker et al., 2019).

Subsequently, variants were called using GATK (v. 4.1.1.0) (McKenna et al., 2010). First, the tool HaplotypeCaller (with the flags --pcr-indel-model NONE and -ploidy 1) was used on each genotype in GVCF mode. Joint genotypes were then produced using the tools CombineGVCFs followed by GenotypeCaller. The raw variants were separated into SNPs and indels and hard-filtered separately according to GATK best practices parameters for quality control; those variants not passing all filters were removed using the SelectVariants tool. The high-quality SNPs and indels were sorted, merged together, and validated using Picardtools (v. 2.18.2) (Wysoker et al., 2019). The variants were functionally annotated using SnpEff (v. 4.3t) (Cingolani et al., 2012) using sequence ontology and a custom reference library built from the annotated RV16 genotype chloroplast genome. The cutoff to label a variant as upstream or downstream from a gene was set at 200bp.

### Phylogenomic analyses of elms and related genomes

For phylogenetic analyses, the two American elm chloroplast genomes were compared to the genomes of the seven Asian elm species listed above and 12 other related species available from Genbank (Table 1). A multiple sequence alignment (MSA) was produced with MAFFT (v. 7.452) using the FFT-NS-i method (Katoh et al., 2002;Katoh and Standley, 2013). Subsequently all gaps created by the alignment were removed by TrimAI in Phylemon 2.0 (Sanchez et al., 2011). To create the phylogeny, the curated MSA was provided to IQ-TREE (v.1.6.11) (Nguyen et al., 2015). First, we used ModelFinder within IQ-TREE to choose the appropriate model of sequence evolution (Kalyaanamoorthy et al., 2017), as assessed by Bayesian Information Criterion (BIC) support measures among 286 DNA models. The best nucleotide substitution model was deemed to be TVM+F+R2. In this nomenclature, TVM represents the base substitution rates of the Transversion model, in which A-G=C-T, with unequal base frequencies. Because the base frequencies were unequal, they were determined empirically from the data. The rate type R represents “free rate” heterogeneity, which relaxes the assumption of Gamma distributed rates and thus is more accurate for large data sets (Yang, 1995;Soubrier et al., 2012). The two categories of rates, in the format of (proportion of sites, relative rate), were determined to be (0.8045, 0.258) and (0.1955, 4.053).

**Table 1.** List of species and NCBI accession numbers used for comparative genomics and phylogenetic analyses.

| Sample name | NCBI accession number |
| --- | --- |
| *Ulmus americana* (RV16) | MN043961.1 |
| *Ulmus americana* (Am. 57845) | MH324448.3 |
| *Ulmus pumila* | NC_032721.1 |
| *Ulmus davidiana* | NC_032718.1 |
| *Ulmus davidiana* var. *japonica* | KY244083.1 |
| *Ulmus macrocarpa* | NC_032720.1 |
| *Ulmus laciniata* | NC_032719.1 |
| *Ulmus gaussenii* | NC_037840.1 |
| *Ulmus chenmouii* | NC_037758.1 |
| *Morus indica* | NC_008359.1 |
| *Morus notabilis* | NC_027110.1 |
| *Morus mongolica* | NC_025772.2 |
| *Ficus racemosa* | NC_028185.1 |
| *Cannabis sativa* | NC_027223.1 |
| *Juglans hopeiensis* | NC_033894.1 |
| *Juglans cathayensis* | NC_033893.1 |
| *Populus trichocarpa* | NC_009143.1 |
| *Salix interior* | NC_024681.1 |
| *Salix purpurea* | NC_026722.1 |
| *Robinia pseudoacacia* | NC_026684.1 |
| *Nicotiana undulata* | NC_016068.1 |

The best substitution model was then used to infer the best phylogeny using a maximum-likelihood (ML) approach. For branch support analysis, we performed 1,000 replicates for a non-parametric Shimodaira–Hasegawa-like approximate likelihood ratio test (SH-aLRT) (Shimodaira and Hasegawa, 1999;Guindon et al., 2010) as well as 1,000 alignments of ultrafast bootstrapping (Hoang et al., 2018). A 50% majority-rule consensus tree was constructed based on 1,000 bootstrap trees. Graphical representation of gene trees was performed with FigTree v. 1.4.3 (http://tree.bio.ed.ac.uk/software/figtree/).

### Intergenic variability in the American elm chloroplast

Intergenic regions of RV16 and Am. 57845 were aligned with Genious software (Grant and Stothard, 2008). Seven intergenic regions were identified as potentially polymorphic between the two American elm genotypes, and these were used to design primers with Primer3 in batch mode (You et al., 2008). Six additional genotypes maintained in the breeding collection were used to evaluate polymorphism in the intergenic spacers (Table 6). For each individual, all genotype-primer combinations were amplified in separate PCR reactions and their amplification tested on 2% agarose gels. The amplicons for each genotype were pooled and submitted for WideSeq analysis at the genomics core facility at Purdue University. WideSeq involves the construction of an NGS library which then is sequenced in its entirety. Raw data were trimmed with trimmomatic software (Bolger et al., 2014). The clean reads were mapped to the reference with BBMap (Bushnell, 2014). GATK HaplotypeCaller (McKenna et al., 2010) was used to identify SNPs and to obtain a depth of coverage file. BCFtools (Narasimhan et al., 2016) was used to generate a consensus sequence, and ClustalW (Larkin et al., 2007) was used to generate an alignment for verifying polymorphic regions.

## RESULTS

### Chloroplast genome organization of American elm

The complete chloroplast genome maps of the American elm genotypes were produced for RV16 (Figure 1) and Am. 57845 (Supplementary Figure 1). The global pairwise nucleotide sequence alignment between the full chloroplast genomes of the RV16 and Am. 57845 genotypes contained 159,106 positions (Supplementary Figure 2). The sequences showed 99.8% identity (158,778/159,106) and 99.8% similarity (158,777/159,106). There were 284/159,106 gap positions (0.2%). The chloroplast genome and region sizes of *U. americana* were similar to those of the seven Asian elm species and closest in size to *U. laciniata* (Table 2). The *U. Americana* complete chloroplast genome assemblies ranged from 158,935bp (Am. 57845) to 158,993bp (RV16). The IR length ranged from 25,563bp (Am. 57845) to 26,072bp (RV16). The SSC ranged from 88,112bp (RV16) to 89,113bp (Am. 57845), and the LLC ranged from 18,696bp (Am. 57845) to 18,737 (RV16) (Table 2, Figure 2). The two genotypes exhibited the same GC content of 35.6%, and the same gene content: 127 genes, including 85 protein-coding genes, 34 tRNA genes and 8 rRNA genes, and were intermediate in the number of genes when compared to the Asian elms (which ranged from 121 in *U. chenmoui* to 145 in *U. davidiana var. japonica*) (Table 2, Table 3). American elm had 16 genes with introns, while the number of genes with introns varied from 8-18 in Asian elms (Table 2).

**Table 2.** Comparison of *Ulmus americana* chloroplast genomes with the published genomes of seven Asian elms.

| Chloroplast<br>Feature | *U. americana*<br>(RV16) | *U. americana*<br>(Am. 57845) | *U.*<br>*pumila* | *U. davidiana*<br>var. *japonica* | *U.*<br>*macrocarpa* | *U.*<br>*davidiana* | *U.*<br>*laciniata* | *U.*<br>*chenmoui* | *U.*<br>*gaussenii* |
| --- | --- | --- | --- | --- | --- | --- | --- | --- | --- |
| Genome size<br>(bp) | 158993 | 158935 | 159082 | 159411 | 159439 | 159453 | 158953 | 159777 | 160094 |
| GC content<br>(%) | 35.6 | 35.6 | 35.6 | 35.6 | 35.6 | 35.6 | 35.6 | 35.5 | 35.5 |
| LSC length<br>(bp) | 88112 | 89113 | 88103 | 88508 | 88647 | 87633 | 88032 | 88581 | 88875 |
| SSC length<br>(bp) | 18737 | 18696 | 18850 | 18868 | 18853 | 18835 | 18846 | 19052 | 19179 |
| IR length<br>(bp) | 26072 | 25563 | 26064 | 26017 | 26019 | 26492 | 26037 | 26072 | 26062 |
| Genes (#) | 127 | 127 | 137 | 145 | 142 | 143 | 139 | 121 | 122 |
| tRNA (#) | 34 | 34 | 46 | 48 | 46 | 47 | 46 | 30 | 30 |
| rRNA (#) | 8 | 8 | 8 | 8 | 8 | 8 | 8 | 8 | 8 |
| Genes with<br>introns (#) | 16 | 16 | 13 | 16 | 16 | 18 | 14 | 8 | 8 |
| Proteins (#) | 84 | 84 | 83 | 89 | 88 | 88 | 85 | 79 | 79 |

**Table 3.**
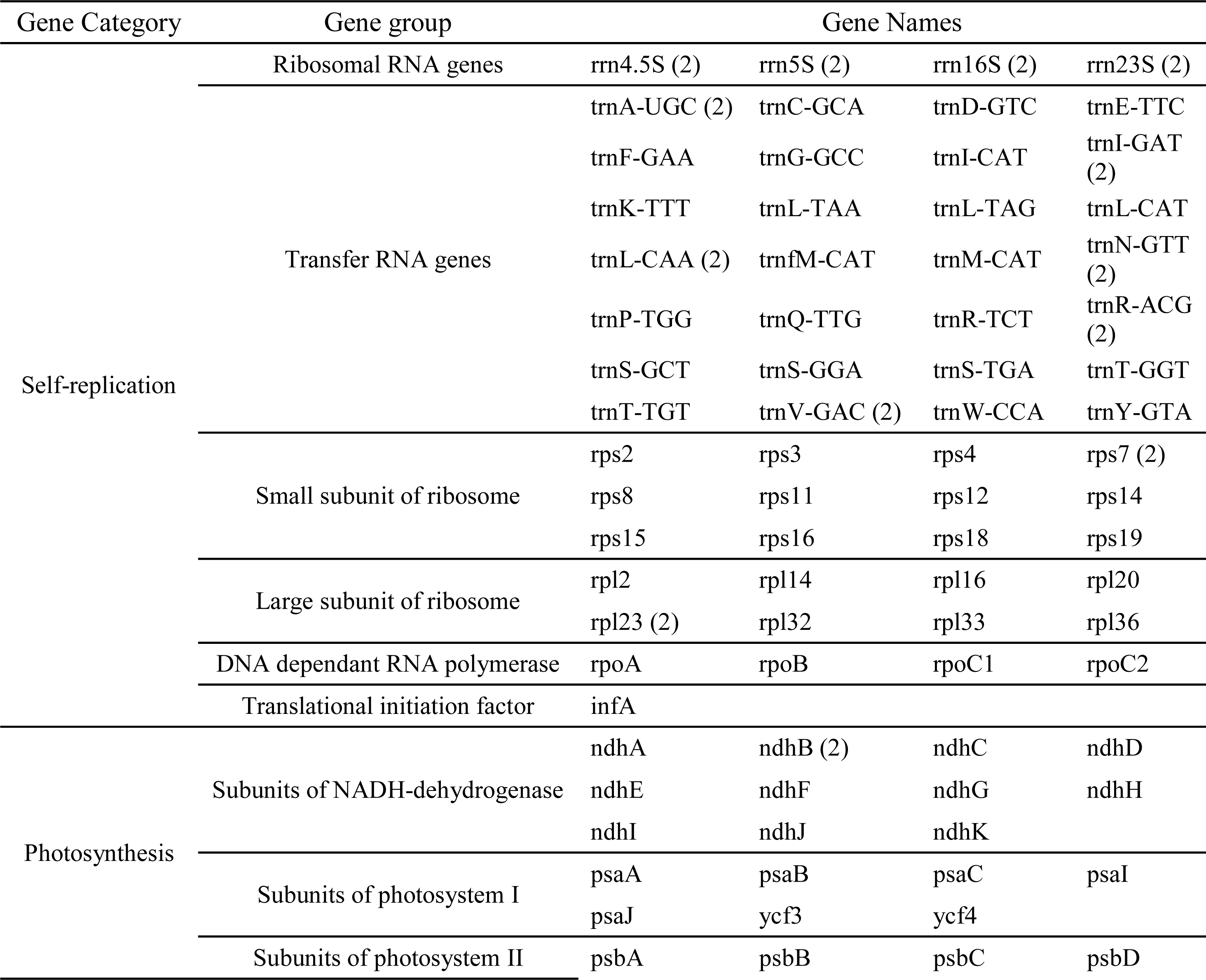
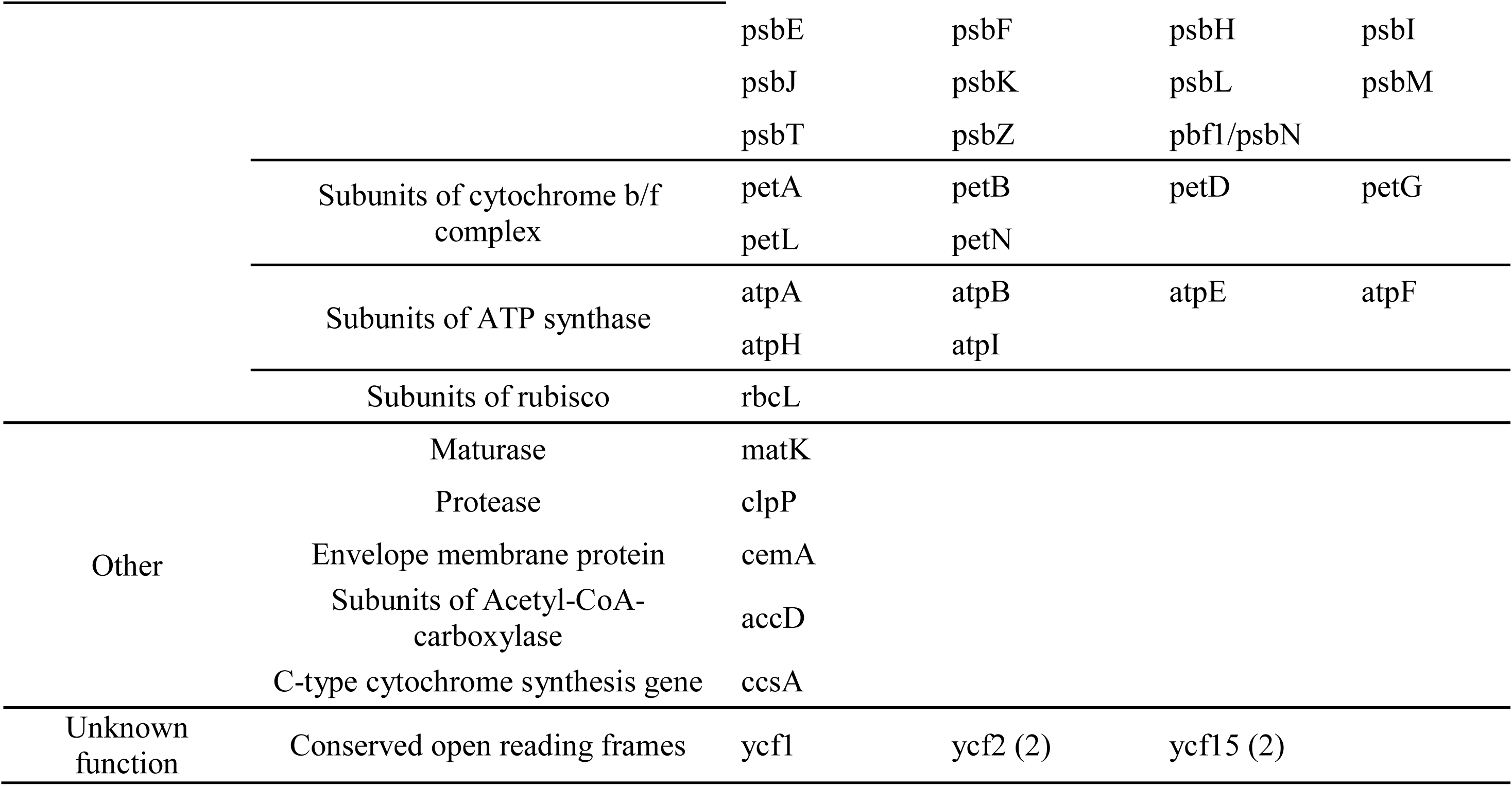
Gene content of the *Ulmus americana* chloroplast genome. Genes with multiple copies are indicated in parentheses.

### Variant identification within the American elm chloroplast

We found *U. americana* chloroplast genome variants in the form of SNPs and indels, which are available in vcf format as Supplementary Table 1. We retained 240 high-quality variants: 67 SNPs, 85 insertions, and 88 deletions (Table 5). Variants occurred at a rate of one per 662bp, and were scattered throughout the chloroplast genome, with the highest number (22) occurring at the 9,000-10,000bp and 124,000-125,000bp ranges of the RV16 genome assembly (Figure 3). These peaks are due to intragenic variants in *trn*L-TAA and intron variants in *ndh*A, respectively (Supplementary Table 1). Upon annotation, we found 328 effects among the 240 variants (as individual variants can exhibit more than one effect) (Table 5). While most variants did not occur in coding regions, variant effects were found acting on 47 different protein coding genes (Supplementary Table 2). Variants deemed high impact occurred in four genes: frame-shift variants in *rpl*2 and *ycf*1 and stop-gained variants in *ndh*F and *ycf*4. Additionally, missense variants (causing amino acid substitutions) were found in *ndh*C, *ndh*K, *rbc*L, *rpl*2, *rpo*C2, and *rps*3 (Supplementary Table 2).

### Gene loss-gain based on the whole chloroplast genome

We evaluated the chloroplast genomes of the eight *Ulmus* species, two species closely related to *Ulmus* (*M. mongolica* and *C. sativa*), and *A. thaliana* for the presence and absence of particular genes (Table 4). Five genes (*pet*B*, pet*D*, psb*L*, rps*16, and *trn*K) were present in American elm but absent from all Asian elms (Zuo et al., 2017;Zhang et al., 2019a;Zhang et al., 2019b). The gene *ycf*15 was present in all elms but absent from the related species *Morus mongolica*, *Cannibis sativa*, and *Arabidopsis thaliana*. The gene *trn*H was present in most Asian elms and the other related species but not in *U. americana*. The gene *trnf*M was present in *U. americana* and the other related species, but absent from over half of the Asian elms (Table 4). Additionally, *U. americana* exhibited only one copy of *ycf*1 (at the junction of SSC and IRb) while all the Asian elms exhibited two, with another copy at the junction of SSC and IRa (Figure 2). However, *U. americana* contained two copies of *rpl*2, one at the junction of LSC and IRa and one at the junction of LSC and IRb, while all Asian elms except *U. davidiana* contained just the copy between the junction of LSC and IRb (Figure 2).

**Table 4.** Presence and absence of certain chloroplast genes in American elm compared with seven Asian elms and species of *Morus*, *Cannabis*, and *Arabidopsis*.

| Species | *petB* | *petD* | *psbL* | *rps16* | *trnK* | *ycf15* | *trnH* | *trnfM* |
| --- | --- | --- | --- | --- | --- | --- | --- | --- |
| *U. americana* | + | + | + | + | + | + | -- | + |
| *U. davidiana* | -- | -- | -- | -- | -- | + | -- | -- |
| *U. davidiana* var. *japonica* | -- | -- | -- | -- | -- | + | + | + |
| *U. pumila* | -- | -- | -- | -- | -- | + | + | -- |
| *U. laciniata* | -- | -- | -- | -- | -- | + | + | -- |
| *U. macrocarpa* | -- | -- | -- | -- | -- | + | + | -- |
| *U. chenmoui* | -- | -- | -- | -- | -- | + | + | + |
| *U. gaussenii* | -- | -- | -- | -- | -- | + | -- | + |
| *M. mongolica* | + | + | + | + | + | -- | + | + |
| *C. sativa* | + | + | + | + | + | -- | + | + |
| *A. thaliana* (NC_000932.1) | + | + | + | + | + | -- | + | + |

**Table 5.** Summary of high-quality variants (SNPs and indels) discovered in the chloroplast of *U. americana* MN043961.1 (RV16) and MH324448.3 (Am. 57845).

| Change rate by chromosome |  |  |
| --- | --- | --- |
| Chromosome | Length | Changes |
| MN043961 | 158993 | 240 |
| Variants by type |  |  |
| Type | Count | Percent |
| Deletion | 88 | 36.7% |
| Insertion | 85 | 35.4% |
| SNP | 67 | 27.9% |
| Effects by impact |  |  |
| Type | Count | Percent |
| High | 4 | 1.2% |
| Low | 6 | 1.8% |
| Moderate | 8 | 2.4% |
| Modifier | 310 | 94.5% |
| Effects by functional class |  |  |
| Type | Count | Percent |
| Missense | 8 | 61.5% |
| Silent | 5 | 38.5% |
| Missense/silent ratio | 1.6 |  |

**Count by effects**
| Type | Count | Percent |
| --- | --- | --- |
| Downstream gene variant | 50 | 15.1% |
| Frameshift variant | 4 | 1.2% |
| Intergenic region | 149 | 44.9% |
| Intragenic variant | 28 | 8.4% |
| Intron variant | 52 | 15.7% |
| Missense variant | 8 | 2.4% |
| Splice region variant | 2 | 0.6% |
| Stop gained | 2 | 0.6% |
| Stop retained variant | 1 | 0.3% |
| Synonymous variant | 4 | 1.2% |
| Upstream gene variant | 32 | 9.6% |

**Count by genomic region**
| Type | Count | Percent |
| --- | --- | --- |
| Downstream | 50 | 15.2% |
| Exon | 16 | 4.9% |
| Intergenic | 149 | 45.4% |
| Intron | 51 | 15.5% |
| Splice site region | 2 | 0.6% |
| Transcript | 28 | 8.5% |
| Upstream | 32 | 9.8% |

**Table 6.**
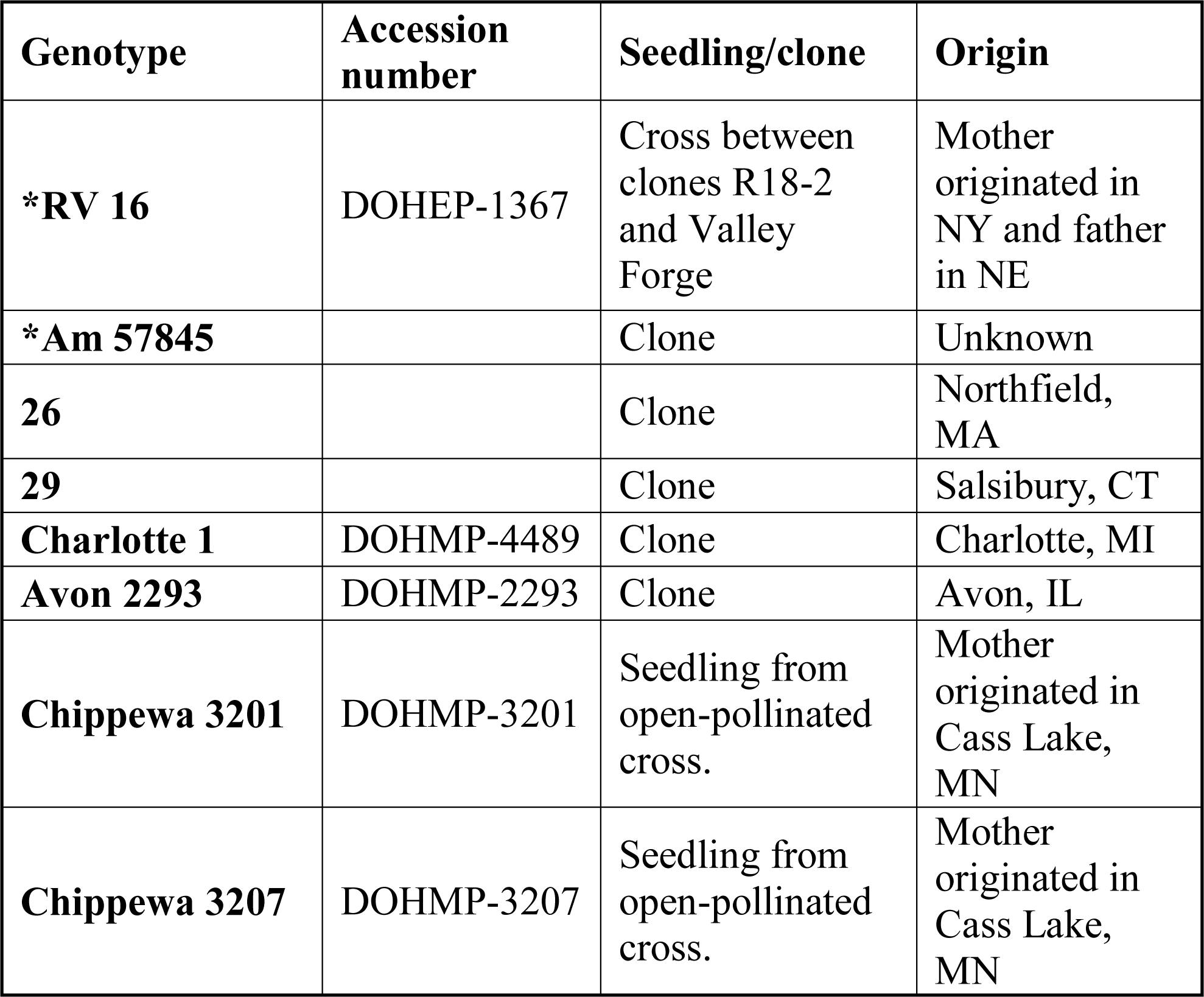
Genotype and origin of the American elm samples used to identify intergenic chloroplast variability. Genotypes with asterisks were used for chloroplast genome assembly, annotation, and additional analyses.

We generated a global multiple sequence alignment of the eight elm species using RV16 (American elm) as the reference sequence for alignment and annotation (Figure 4 and Supplementary Figure 3). These identity plots show that Asian elms, while similar among themselves, differ from American elm, with non-coding regions less conserved within the genus than exons.

### Phylogenomic analyses

We prepared a phylogeny based on the chloroplast genomes of 21 species that ranged from 153,854-160,094bp, and produced an initial MSA of 198,896 positions. Upon gap removal, 119,518 nucleotide positions remained, of which 17,831 were parsimony-informative and 76.2% were constant.

Using the MSA and best model of nucleotide substitution, the best maximum likelihood tree had a total tree length (sum of branch lengths) of 0.4272. The sum of internal branch lengths was 0.1936 (45.3% of the total tree length). The majority rule consensus tree based on 1000 ML bootstrap trees had the same topology as the best ML tree. The consensus tree represents the phylogeny for the 21 species, pictured as both a rectangular phylogram (in which branch lengths represent inferred evolutionary change as the number of nucleotide substitutions per site) and as a cladogram (with equal branch lengths) (Fig. 5). The American elms clustered separately from the Asian elms, and all elms sorted in a clade distinct from the other species included.

### Intergenic variability in the American elm chloroplast

We identified seven intergenic regions that appeared polymorphic between the two American elm chloroplast genomes. Based on resequencing in these regions (27,988 reads with > 10X coverage for each putative polymorphism) among six additional American elm samples (Table 6), alignment of the resequencing reads, and SNP calling, we found that the *rps*12*-ycf*15 intergenic spacer contained 84 SNPs among the six genotypes, the *mat*K-*trn*K-UUU spacer contained 37 SNPs, the *trn*K-UUU*-rps*16 spacer contained 9 SNPs, and the *trn*M-CAU*-psb*D showed 8 SNPs.

## DISCUSSION

We assembled from paired-end whole genome sequence data the whole chloroplast genomes of two American elm genotypes. The availability of these sequence data and identified variants increases considerably our knowledge of the chloroplast of American elm and provides a resource for American elm population genetics. Chloroplast variation can be used to understand genetic variation, spatial genetic structure, and gene flow, and is important for taxonomy (Wiegrefe et al., 1994). Sequence variation in uniparentally-inherited chloroplasts can also be used to detect ancestral and recent hybridization of American elm with other elm species, a common feature of modern elm breeding. Most *U. americana* are tetraploid (4n=56), but Whittemore and Olsen (2011) reported 20 percent of American elms were also diploid (2n=28). The presence of diploid *U. americana* opens the possibility of gene exchange between American elm and other diploid elm species, e.g., the exotic *U. pumila* (Brunet and Guries, 2016). Hybridization between American elm and the native sympatric diploid *U. rubra* seems improbable and has never been reported.

### Chloroplast genome organization of American elm and comparison with Asian elm

The genome of American elm was intermediate in size in comparison with the Asian elms. A comparison of American elm chloroplasts with Asian elm chloroplast genomes showed variability in gene content, number of coding genes, and genome size (Table 2). For example, the *pet*B*, pet*D*, psb*L*, rps*16, and *trn*K were identified in the chloroplasts of *U. americana* but absent from the chloroplasts of Asian elm species (Table 4). *trn*H, which encodes a histidine-accepting tRNA, is absent from the chloroplast of *U. americana* but present in the chloroplast of most Asian elms and related species (Table 4). Angiosperms show considerable variation in how they manage tRNAs missing from chloroplasts; they can be expressed from either the mitochondria or from the nucleus and then imported into the mitochondria (Salinas-Giegé et al., 2015).

The genes *pet*B and *pet*D were present in American elm but were lost in the chloroplasts of all Asian elms (Table 4). In plants, *pet*B and *pet*D are members of the *psb*B operon (Soltis et al., 1998), and the *pet*D gene is necessary for photosynthesis (Huang et al., 2017). The *psb*B gene, which has been well studied in higher plants (Stoppel and Meurer, 2013) along with *psb*T and *psb*H have an important roles in photosystem II (PSII). *pet*B is a component of cytochrome b6, which facilitates electron transfer between photosystem II and photosystem I (Stoppel and Meurer, 2013). The absence of *pet*B and *pet*D in Asian elm chloroplasts means those genes are found in the nucleus of those species (Lin et al., 2015). *psb*L, which encodes an essential protein of photosystem II (Rochaix et al., 1989;Cai et al., 2015;Huang et al., 2017), was also present in American elm chloroplasts but absent from the chloroplasts of Asian elm species. The loss or gain of genes from the chloroplast (i.e., their capture by the nuclear genome) has been considered a consequence of adaptive evolution (Huang et al., 2017). If so, the preservation or loss in *Ulmus* of the genes discussed above reflects an aspect of *Ulmus* evolution that is not yet understood. The study of the genetic diversity of the chloroplasts of additional *Ulmus* species across their entire range may clarify the relationship (if any) between gene loss in *Ulmus* and its biogeography. The genetic structure of cytoplasmic haplotypes in diploid versus tetraploid American elms may inform the nature and locations of hybrid zones (if any) between these populations and, at a more basic level, inform studies of how nuclear genome duplication affects the assembly of multi-subunit cytonuclear complexes, a poorly understood topic (Birchler and Veitia, 2012). Uncertainty about the role of chloroplast gene loss in evolution is compounded by homoplasy (e.g., non-sister species may independently shift a gene from the chloroplast to the nucleus) and by a lack of information about the coordination of expression and function of some organellar genes (Huang et al., 2017).

### Variant identification within the American elm chloroplast

We identified and functionally annotated 240 high-quality SNPs and indels in the *U. americana* chloroplast genome. Although we did not experimentally examine the function of each protein-coding gene with an effect from a variant, our analyses allow us to predict certain changes as having a large effect on gene expression or function. The genes with high-impact effects (i.e. frame-shift or stop-gained variants) and/or missense variants (resulting in amino acid changes) are most likely to influence or reflect evolutionary change in American elm. These include genes involved in photosynthesis, namely subunits of NADH-dehydrogenase (*ndh*C, *ndh*F, and *ndh*K), a subunit of Photosystem I (*ycf*4), and a subunit of rubisco (*rbc*L) (Table 3, Supplementary Table 2). High-impact and/or missense variants were also predicted in genes involved in self-replication, including in the large subunit of the ribosome (*rpl*2) and in DNA-dependent RNA polymerase (*rpo*C2), as well as in a conserved open reading frame with unknown function (*ycf*1) (Table 3, Supplementary Table 2). Additionally, the exceptionally large number of variants found intragenically in the transfer RNA *trn*L-TAA and in the intron of *ndh*A, another subunit of NADH-dehydrogenase, may be of biological importance (Figure 3, Supplementary Table 1). Further study of these genes could elucidate what consequences, if any, these changes have for adaptation or development in American elm.

### Phylogeny of Ulmaceae

We compared the whole chloroplast genome of two American elm genotypes with seven Asian elm species, species of *Morus*, *Ficus*, and *Cannabis*, and seven members of related genera as outgroup species (Table 1, Figure 5). American elm sorted closely with the Asian elms in a sister clade with representatives in the *Cannabaceae* and *Moraceae*, as expected (Zuo et al., 2017). The seven Asian elms sorted into subgroup that was not shared by American elm, which diverged earlier (Figure 5). A strong division between the western and eastern elms based on chloroplast and nuclear (rDNA) was previously noted (Wiegrefe et al., 1994). Our phylogeny agreed with the Zuo et al. (2017) study of five Asian elms which showed that the two *U. davidiana* lineages were not monophyletic. Our phylogeny clarified the timing of the divergence of *U. chenmoui* and *U. gaussenii* relative to the other Asian elms (Zhang et al., 2019a;Zhang et al., 2019b) and American elm.

### Intergenic variability in the American elm chloroplast

After identifying putative polymorphic intergenic regions between the RV16 and Am. 57845 genotypes via alignment, we validated the candidate markers by resequencing those regions in six genotypes of American elm (Table 6). Among these, we identified four polymorphic intergenic spacers: *rps*12*-ycf*15, *mat*K-*trn*K-UUU, *trn*K-UUU*-rps*16, and *trn*M-CAU*-psb*D. Further analysis of these intergenic spacers and the above-reported SNPs and indels within the *U. americana* chloroplast should prove useful for understanding American elm genetic diversity and structure.

### Conclusion

We present the first complete chloroplast genome of American elm. The whole chloroplast genome data (deposited as MH324448 and MN043961) will be useful as a reference for assembling and annotating chloroplast genomes of other New World elms. Along with the chloroplast genome variants identified, these data can be used for as a tool for studying genetic diversity of American elm and for evaluating hybridization between American elm and Asian elm species.

## ACKNOWLEGMENTS

Mention of a trademark, proprietary product, or vendor does not constitute a guarantee or warranty of the product by the U.S. Dept. of Agriculture and does not imply its approval to the exclusion of other products or vendors that also may be suitable. DNA sequencing was performed at the Purdue University Agricultural Genomics Center, Philip San Miguel, Director.

## AUTHOR CONTRIBUTION STATEMENT

AE: conceived and designed the project, analyzed the data and wrote the original manuscript; JDA: designed portions of the methods, analyzed the data, and revised and edited the final manuscript; CCP, JMS, CEF: provided leaves samples and revised the draft; KEW: supervised the project, edited and revised the manuscript. All authors contributed the editing of final manuscript.

## CONTRIBUTION TO THE FIELD STATEMENT

Genetic diversity within the chloroplast is considered a fundamental tool for understanding population genetic structure and species evolution. We report the assembly and public deposit of the full chloroplast DNA sequence of two genotypes of American elm. These sequences enabled us to identify highly polymorphic regions within the American elm chloroplast that can be used for future studies of genetic diversity, genetic structure, gene flow, hybridization, and genome evolution of American elm.

## FIGURE CAPTIONS

**Figure 1.** Map of the chloroplast genome of *Ulmus americana* (RV16). The direction of transcription is indicated by arrows. Genes inside the circle are transcribed clockwise, and those outside are transcribed counterclockwise. Gene functions are color-coded as shown in the legend. The darker gray in the inner circle shows the GC content, while the lighter gray shows the AT content. LSC (Large Single Copy region), SSC (Small Single Copy region), IRA, IRB (Inverted Repeat A and B, respectively). The bold black arrow indicates the start position of the chloroplast assembly; the numbering proceeds counter-clockwise.

**Figure 2.** Chloroplast region (IR/SSC/LSC) junctions among American elm and the seven Asian elms. The bold black arrow indicates the start position of the chloroplast assembly; the numbering proceeds to the left.

**Figure 3.** Histogram showing the distribution of variants (SNPs and indels) discovered in *U. americana* chloroplast from Am. 57845 and RV16 sequencing reads, using the RV16 assembly as a reference. Numbering is along the RV16 assembly, with a bin size of 1,000bp.

**Figure 4.** Sequence identities between the eight globally-aligned *Ulmus* genomes using the RV16 genotype of *U. americana* as a reference for alignment and annotation. Genome regions are color-coded according to the legend. Non-protein-coding genes such as tRNAs are not labeled.

**Figure 5:** Phylogeny based on chloroplast sequences of American elm *(Ulmus americana)*, seven Asian elms *(Ulmus spp.)*, Mulberry *(Morus indica; Morus notabilis; Morus mongolica)*, Fig *(Ficus racemosa)*, and Cannabis *(Cannabis sativa)*, genera belonging to the same tribe as elms, and some outgroup species as listed in Table 1. Trees are unrooted but outgroup black locust (*Robinia pseudoacacia*) is drawn at the root. (A) Rectangular phylogram, in which branch lengths represent inferred evolutionary change as the number of nucleotide substitutions per site; (B) The same tree as a cladogram with equal branch lengths.

## CAPTIONS FOR SUPPLEMENTARY FIGURES AND TABLES

**Supplementary Figure 1.** Map of the chloroplast genome of *Ulmus americana* (Am. 57845). The direction of transcription is indicated by arrows. Genes inside the circle are transcribed clockwise, and those outside are transcribed counterclockwise. Genes of different functions are color-coded as shown in the legend. The darker gray in the inner circle shows the GC content, while the lighter gray shows the AT content. LSC (Large Single Copy region), SSC (Small Single Copy region), IRA, IRB (Inverted Repeat A and B, respectively). The bold black arrow indicates the start position of the chloroplast assembly; the numbering proceeds counter-clockwise.

**Supplementary Figure 2.** Pairwise global nucleotide sequence alignment of *Ulmus americana* genotypes RV16 and Am. 57845.

**Supplementary Figure 3.** Larger version of Figure 4. Sequence identities between the eight *Ulmus* genomes using the RV16 genotype of *U. americana* as a reference for alignment and annotation. Genome regions are color-coded according to the legend. Non-protein-coding genes such as tRNAs are not labeled.

**Supplementary Table 1.** The vcf file containing all high-quality variants (SNPs and indels) discovered in the *U. americana* chloroplast, including functional annotations.

**Supplementary Table 2.** The genes in the *U. americana* chloroplast affected by variants (SNPs and indels).

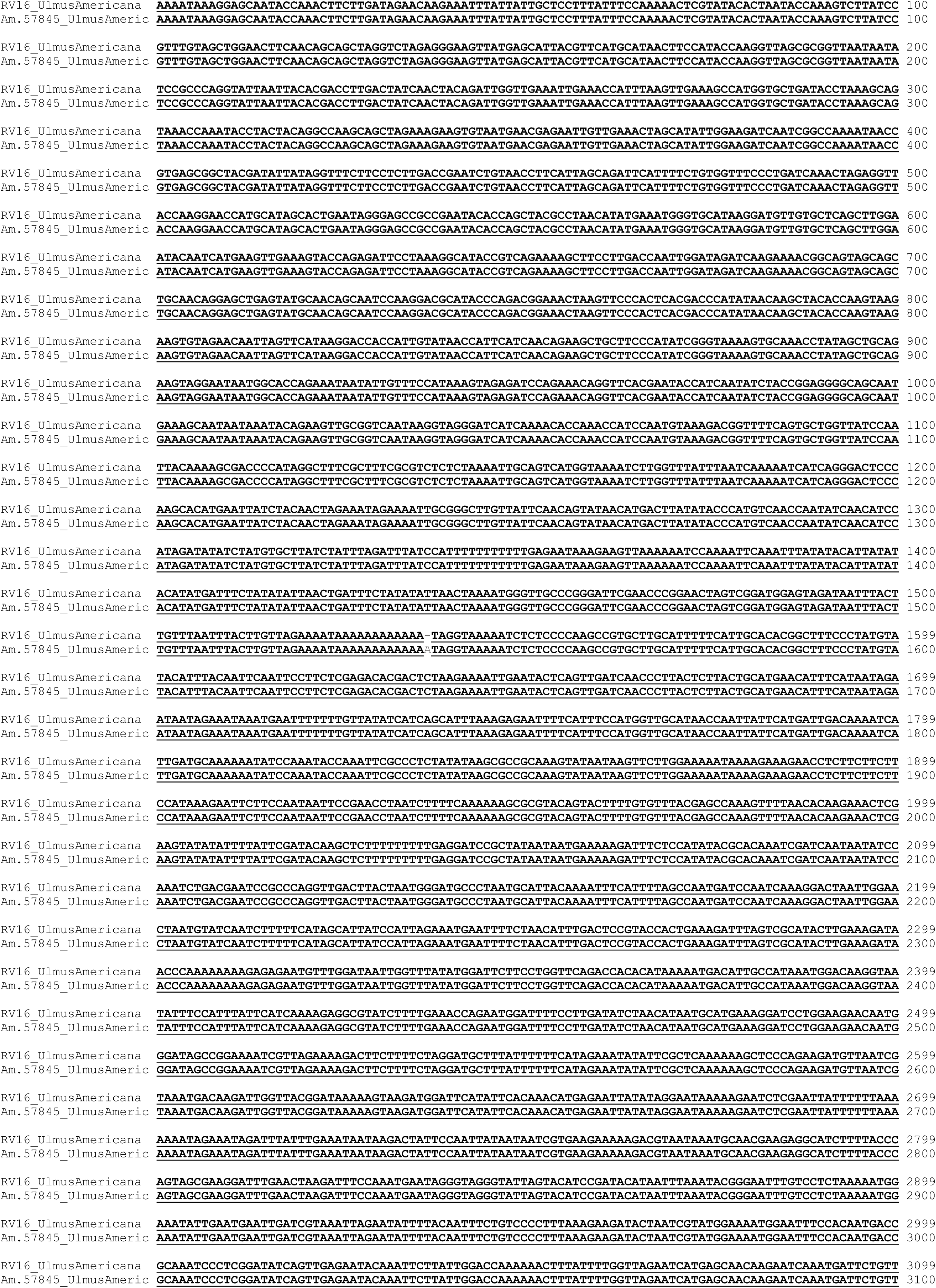
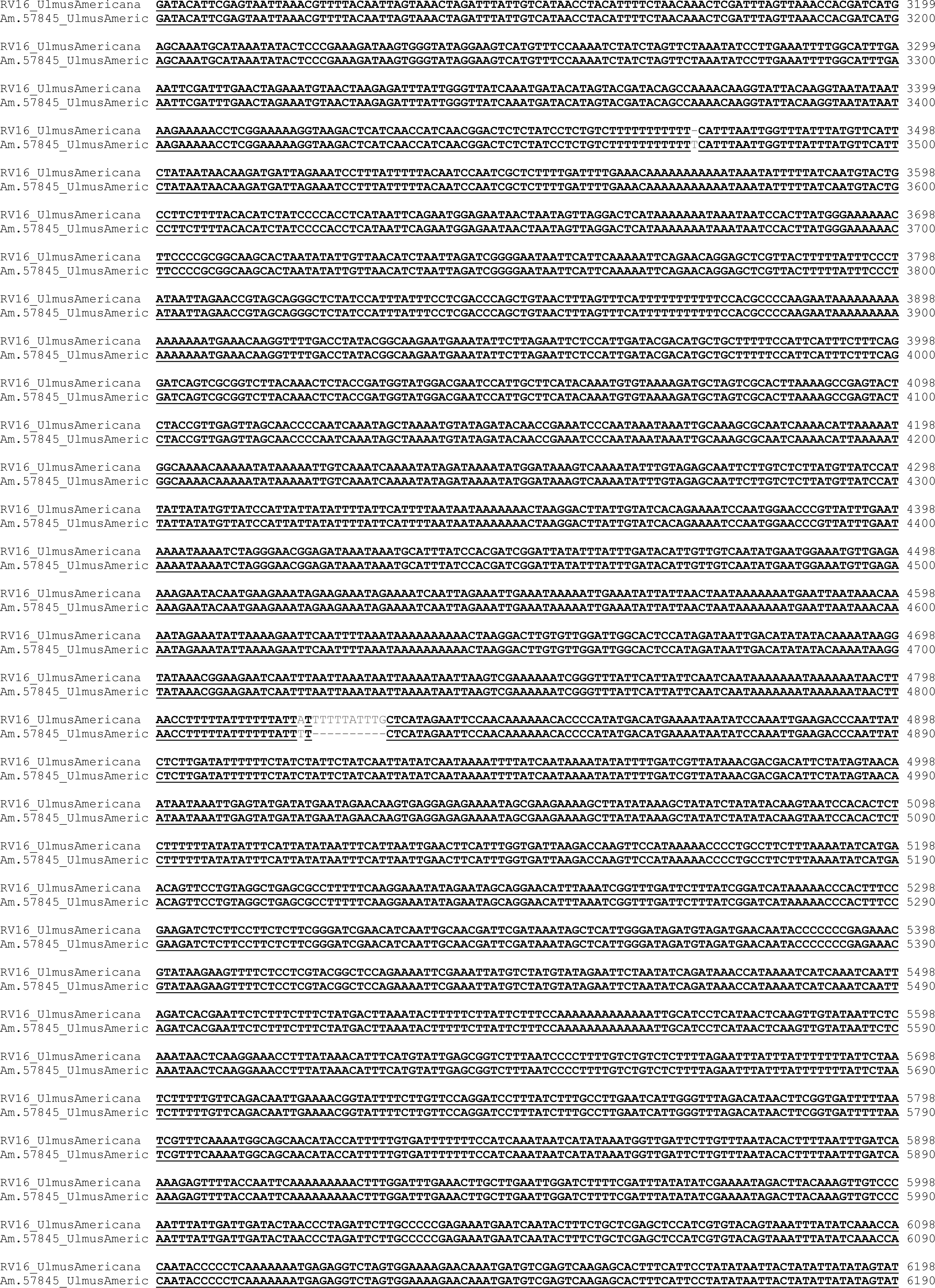
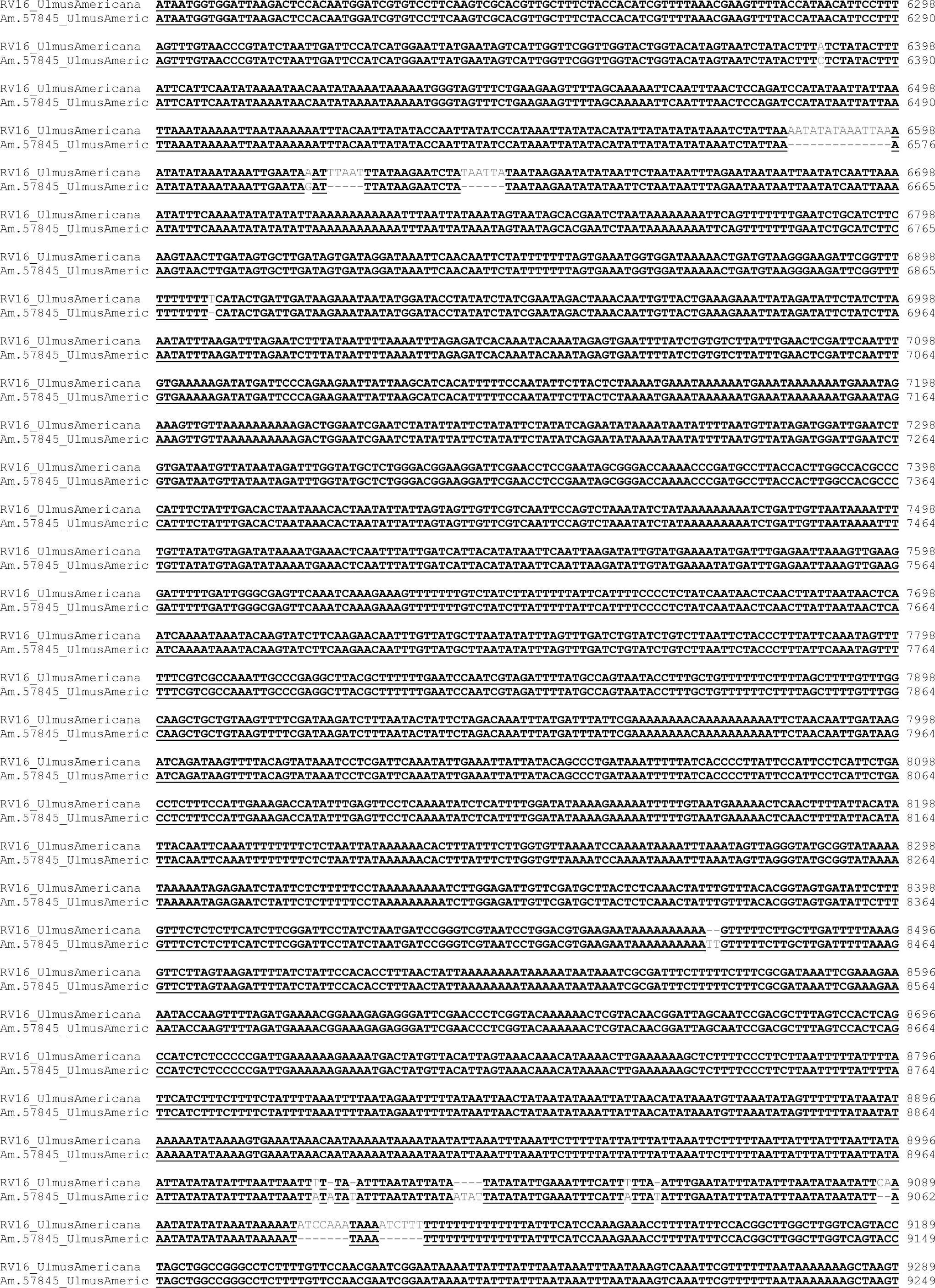
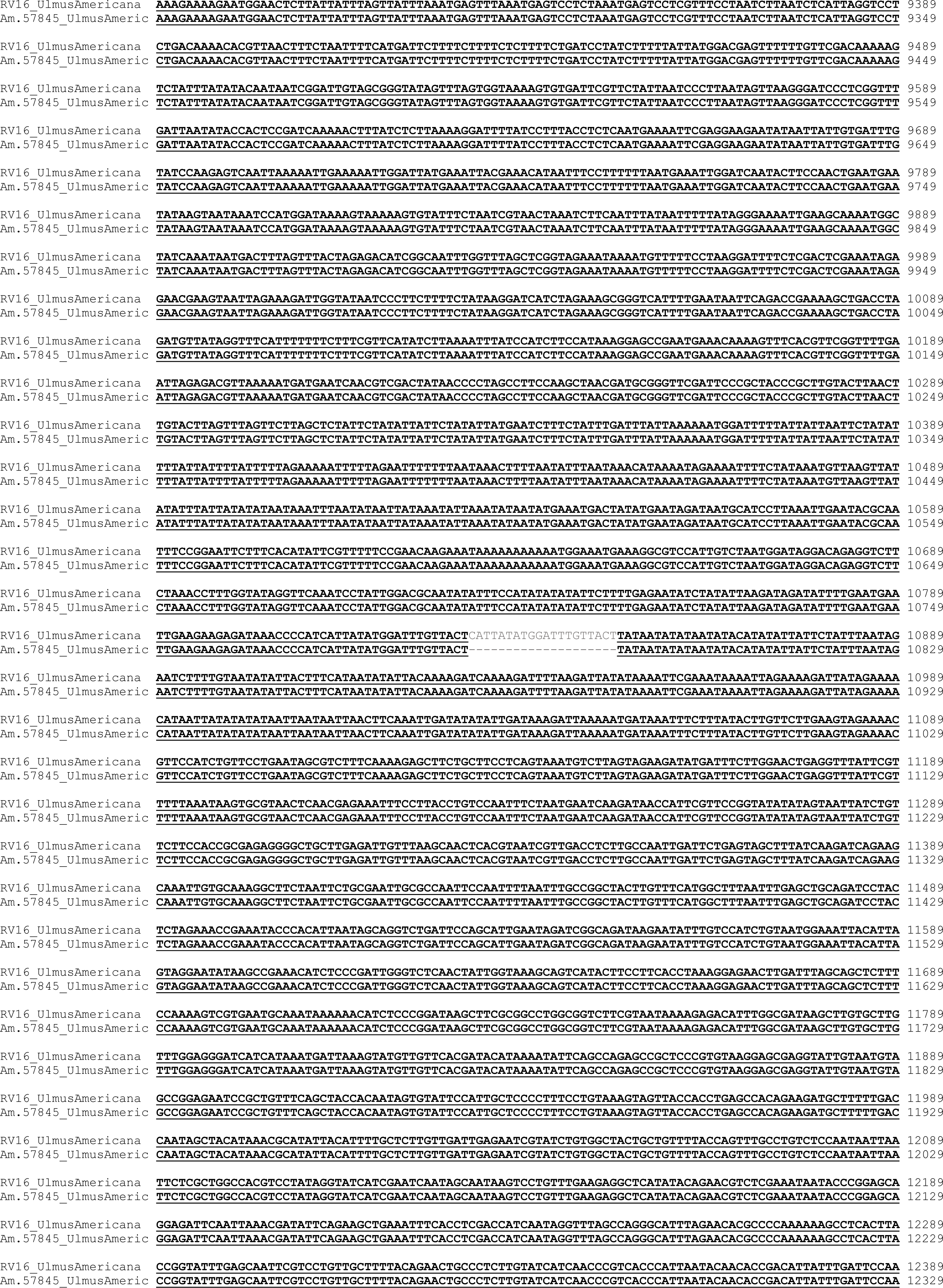
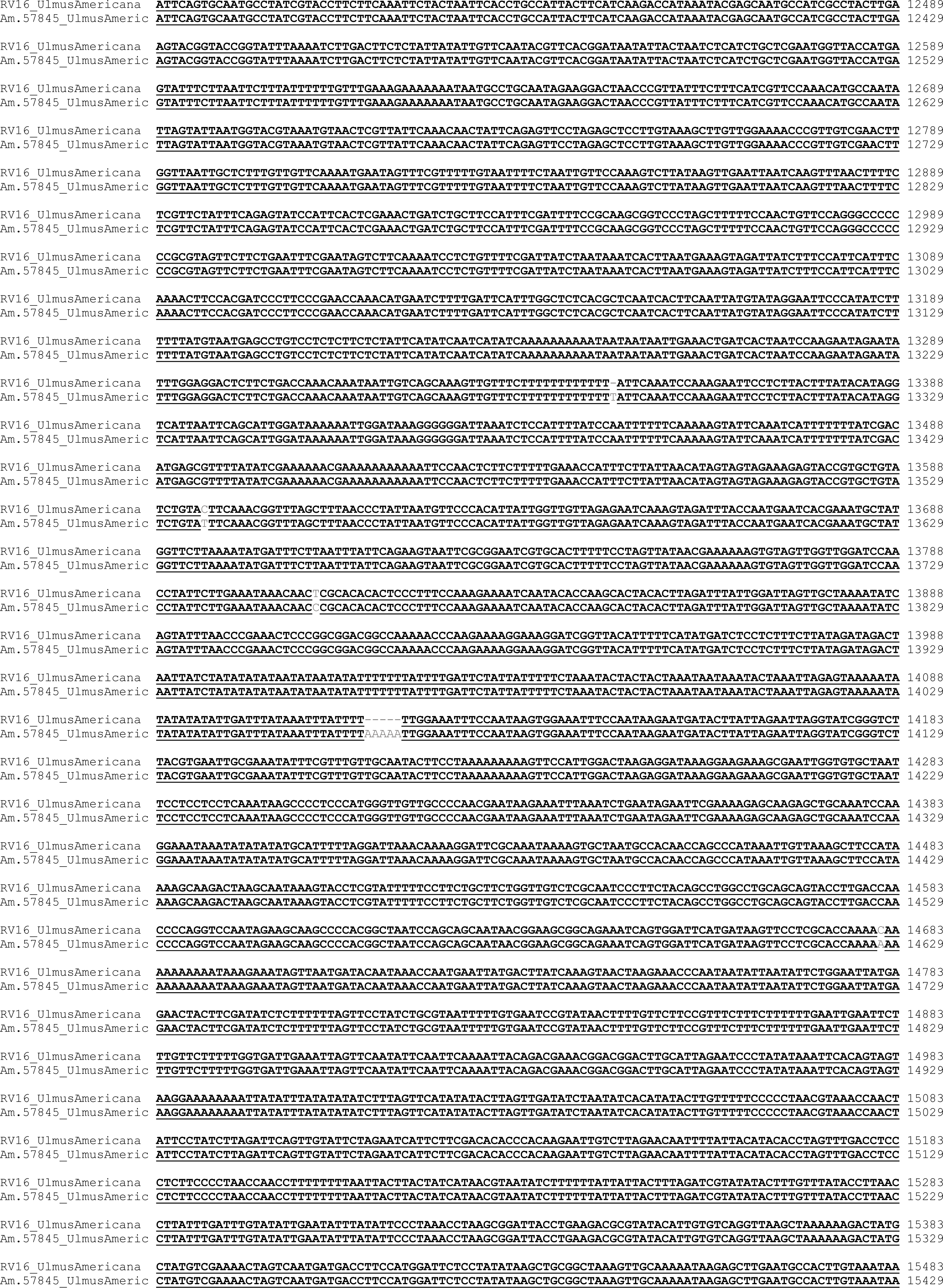
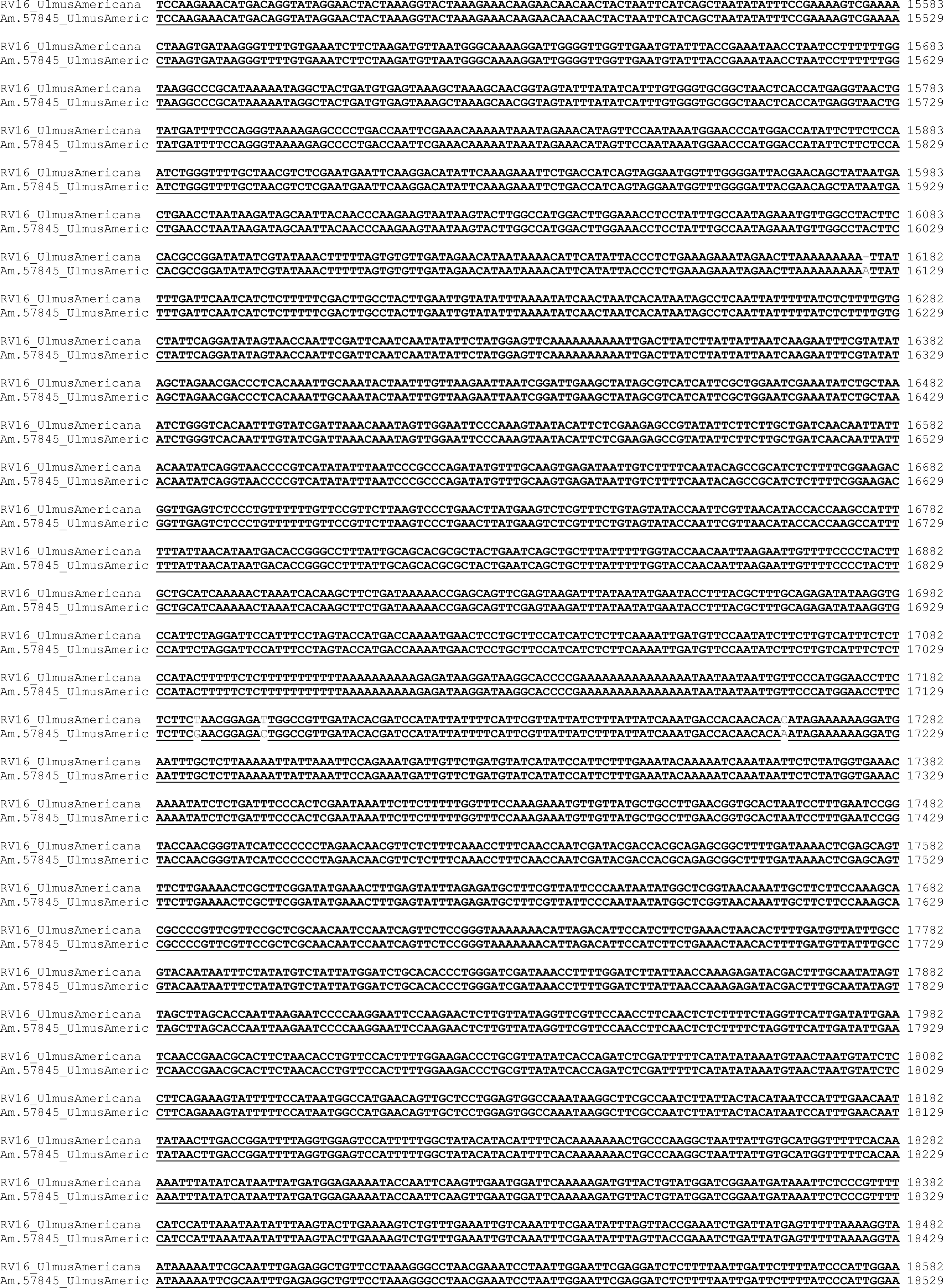
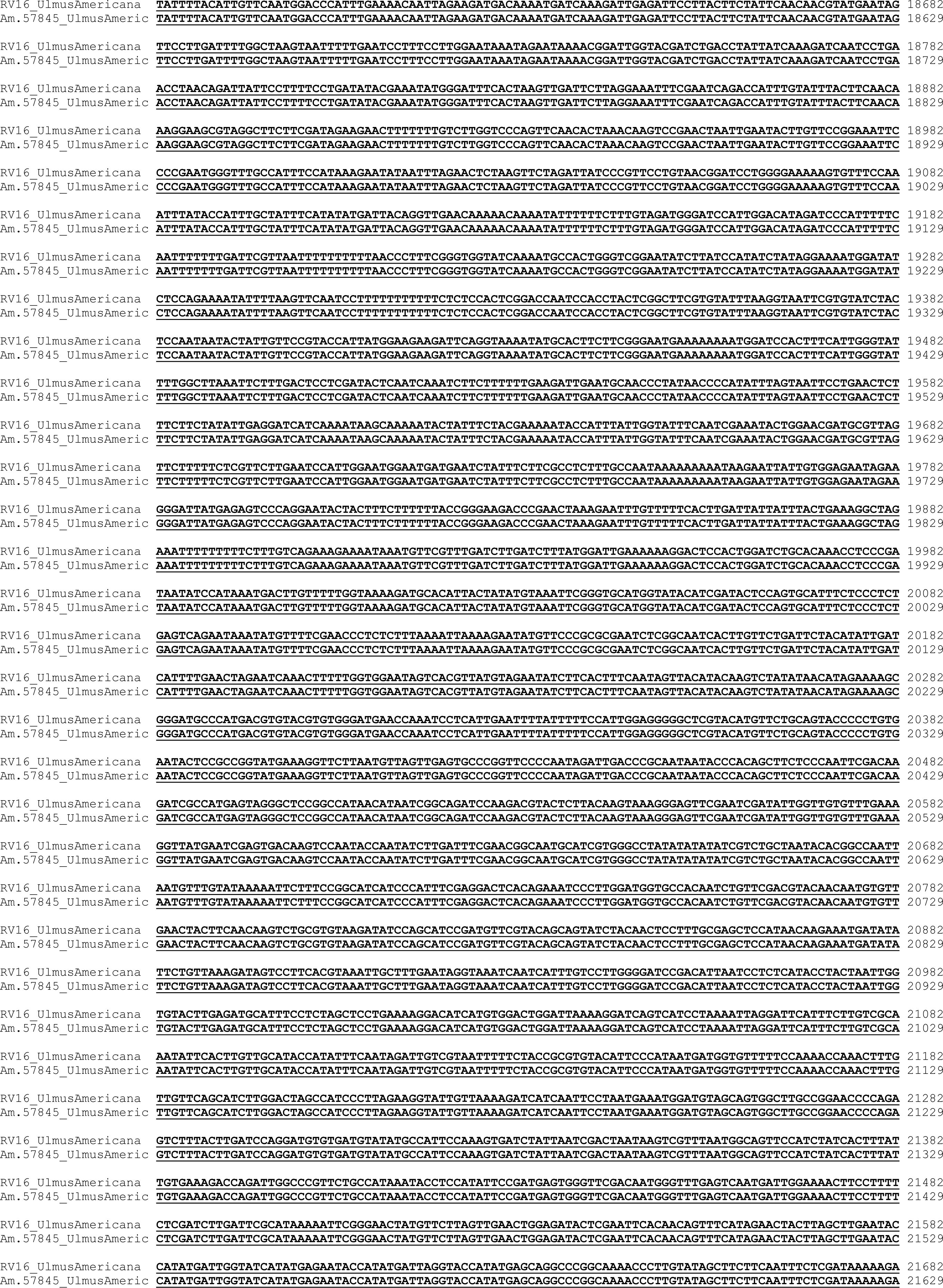
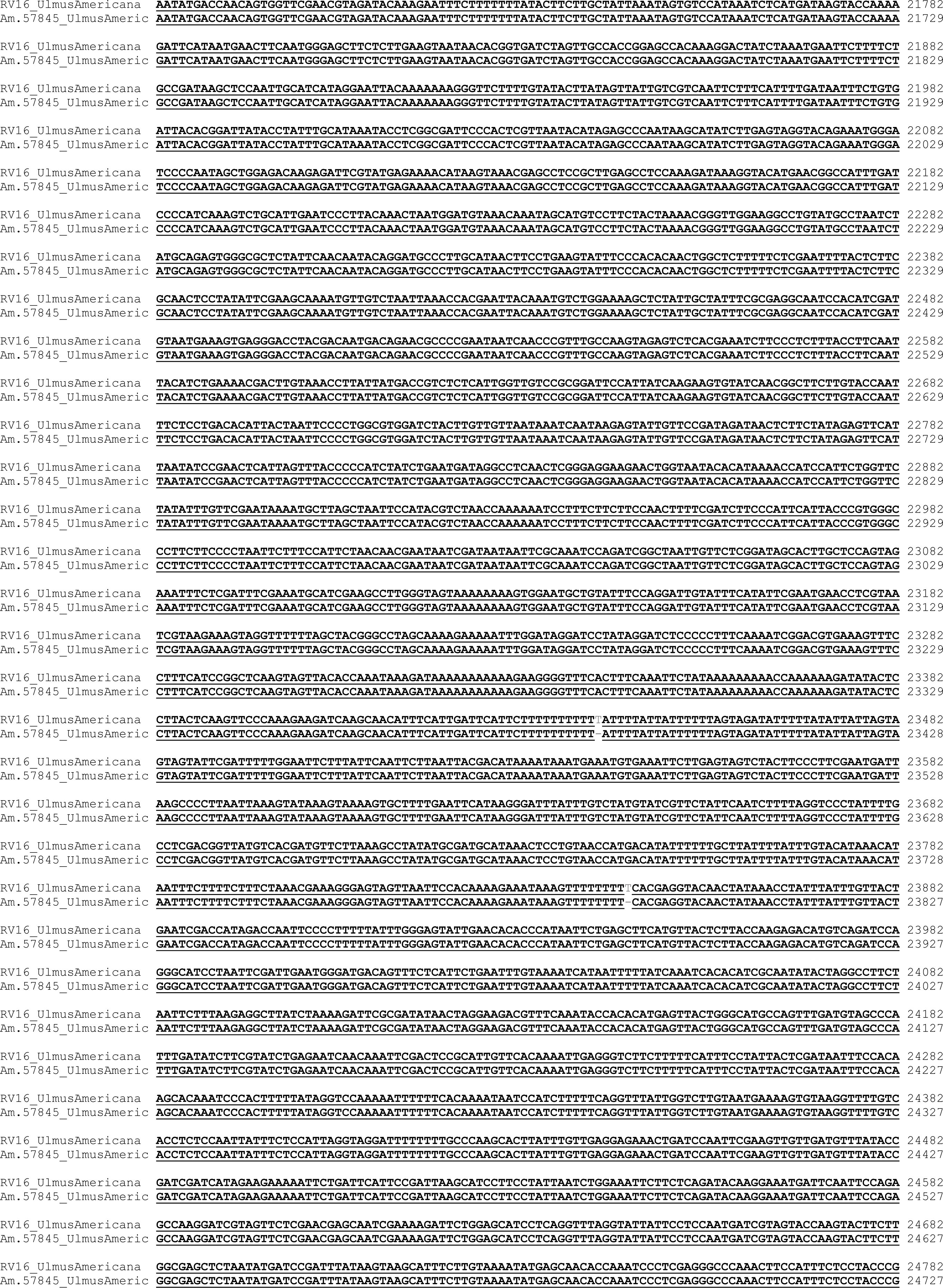
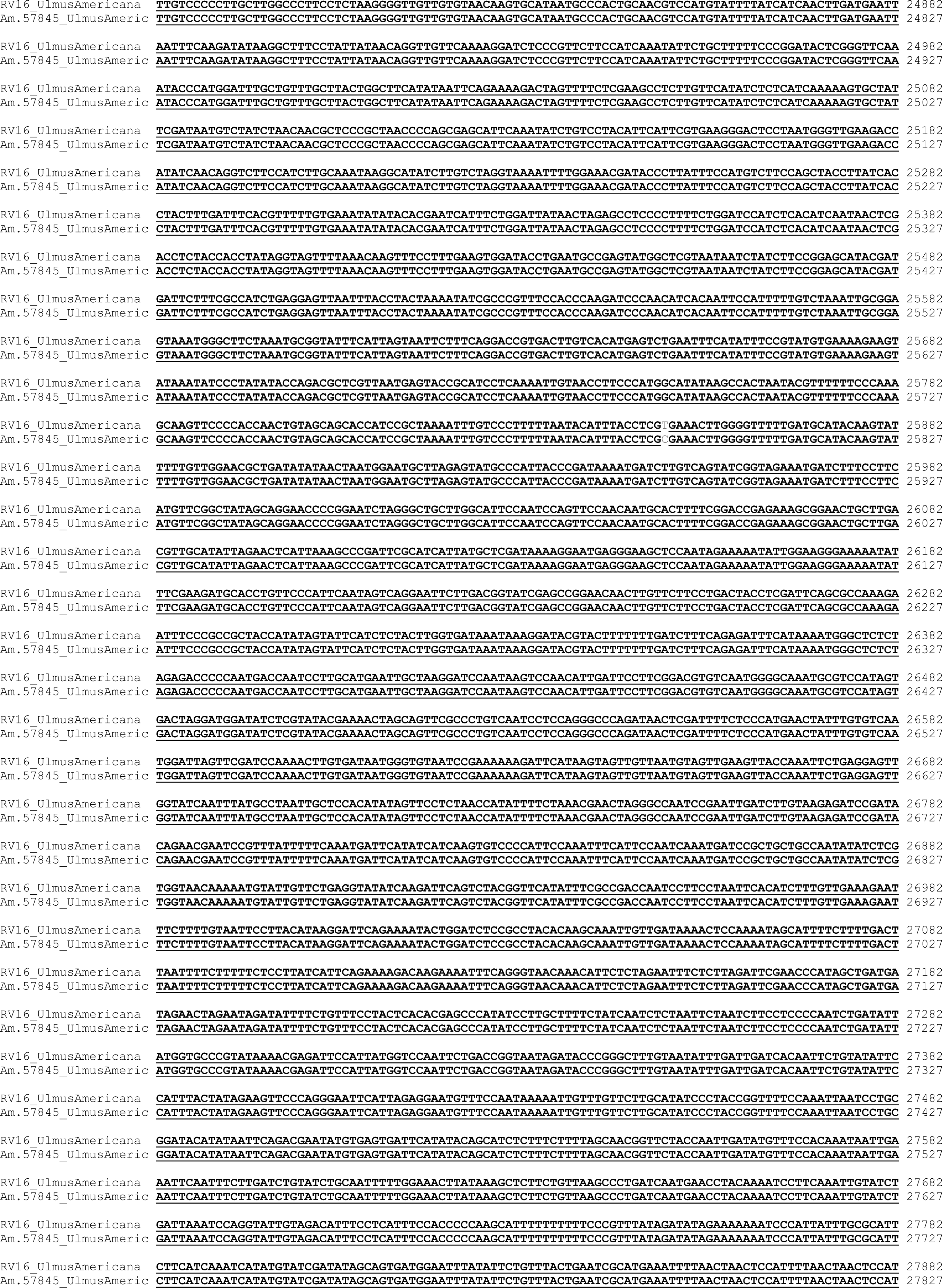
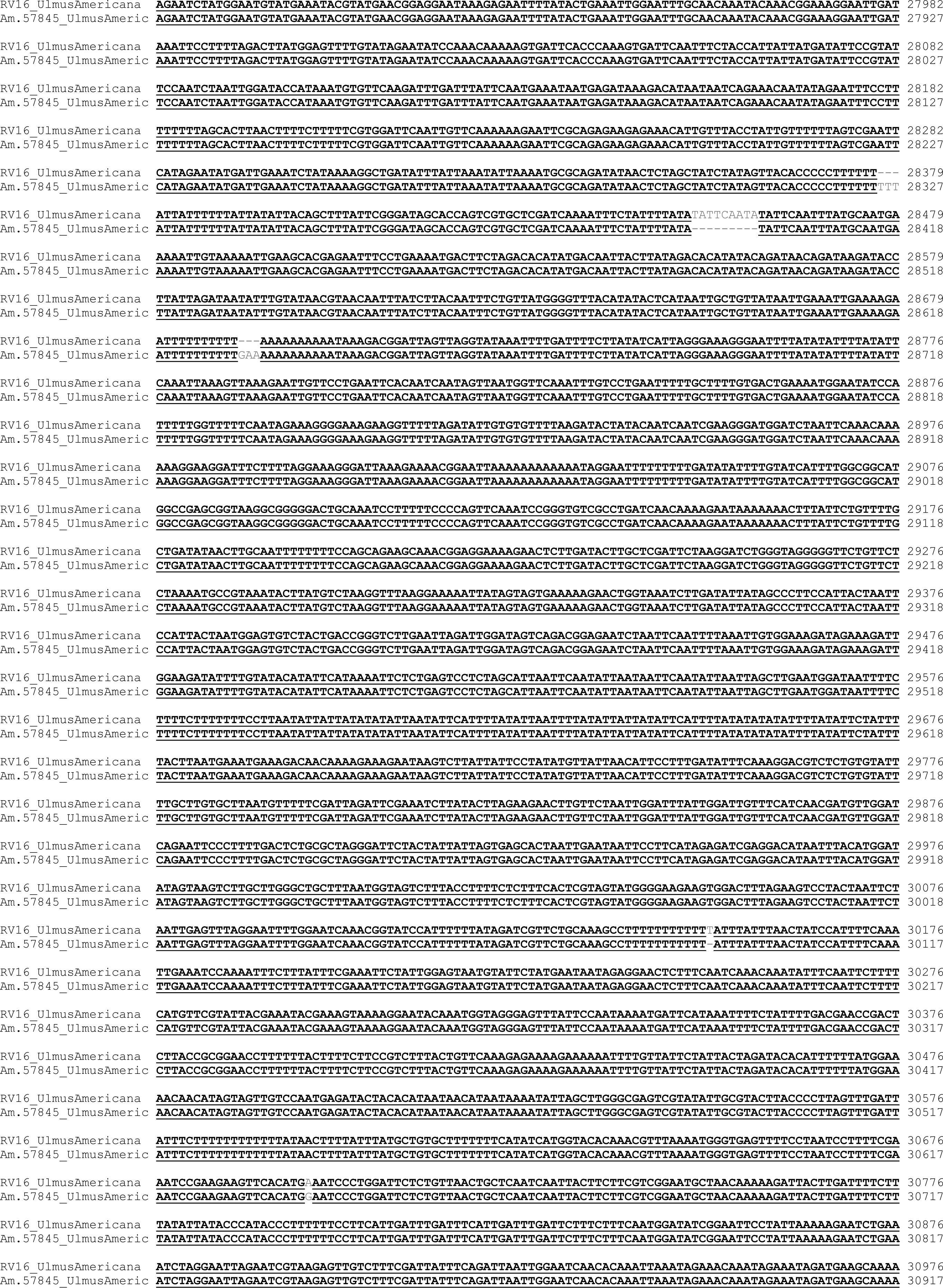
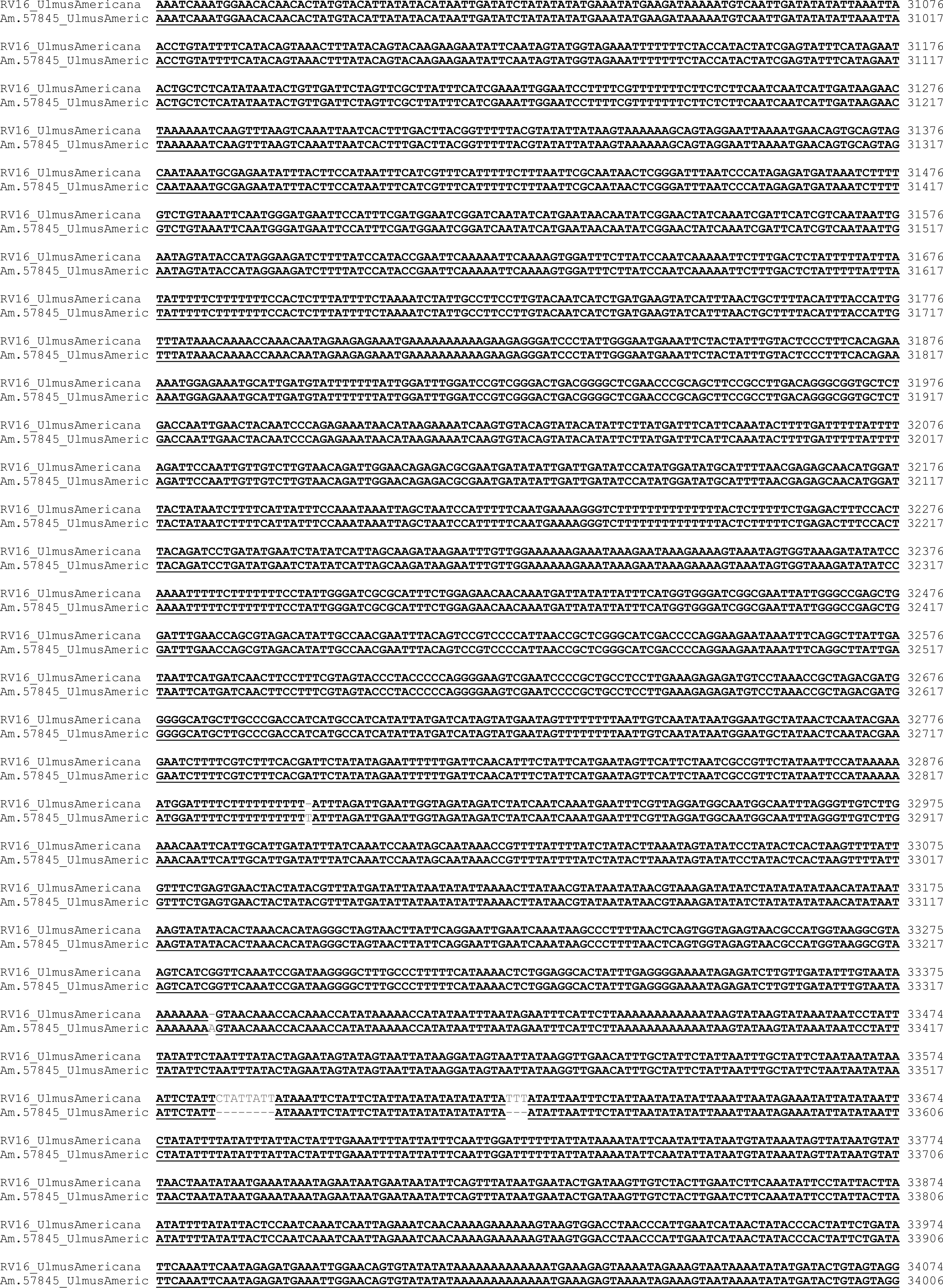
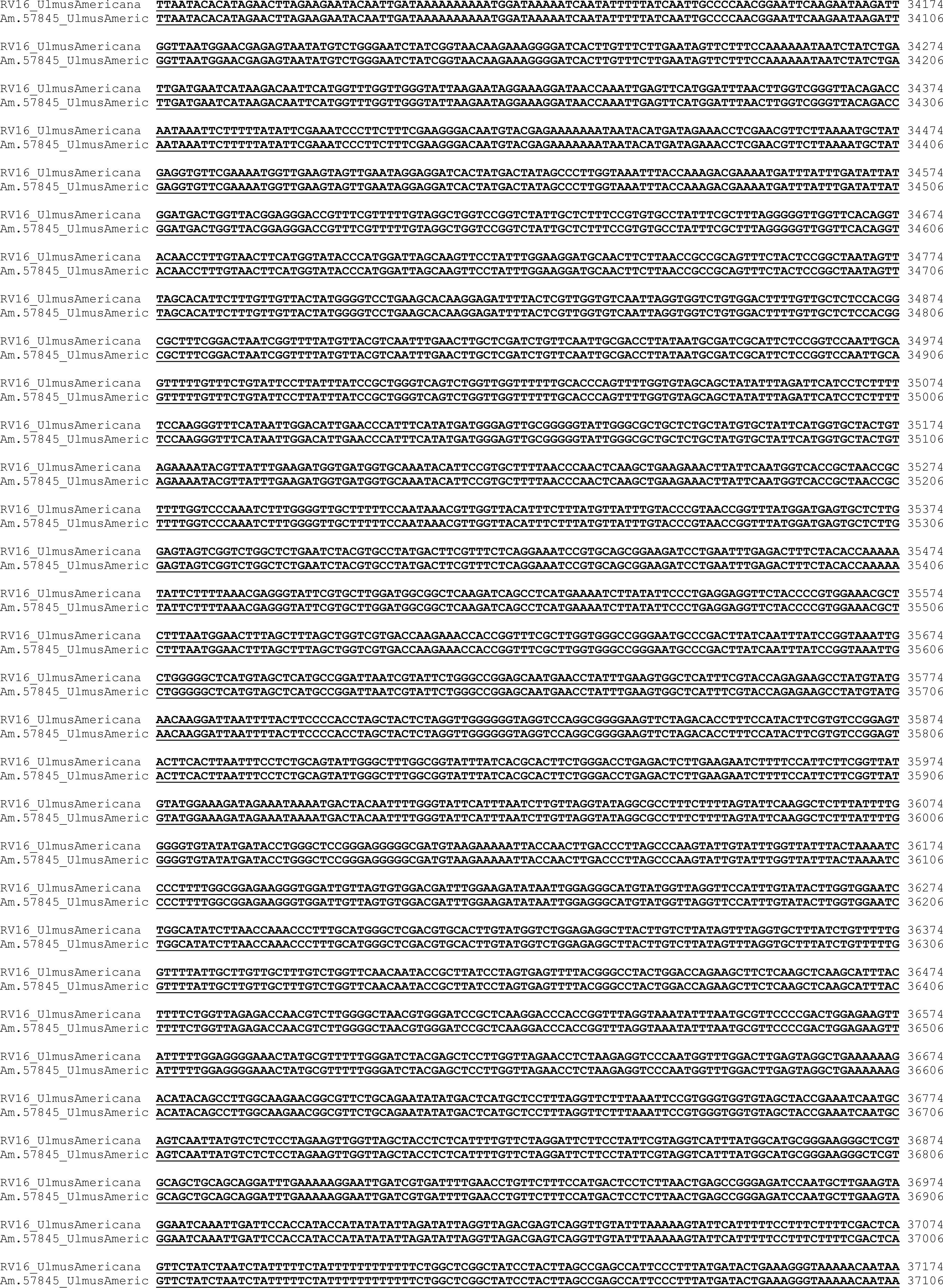
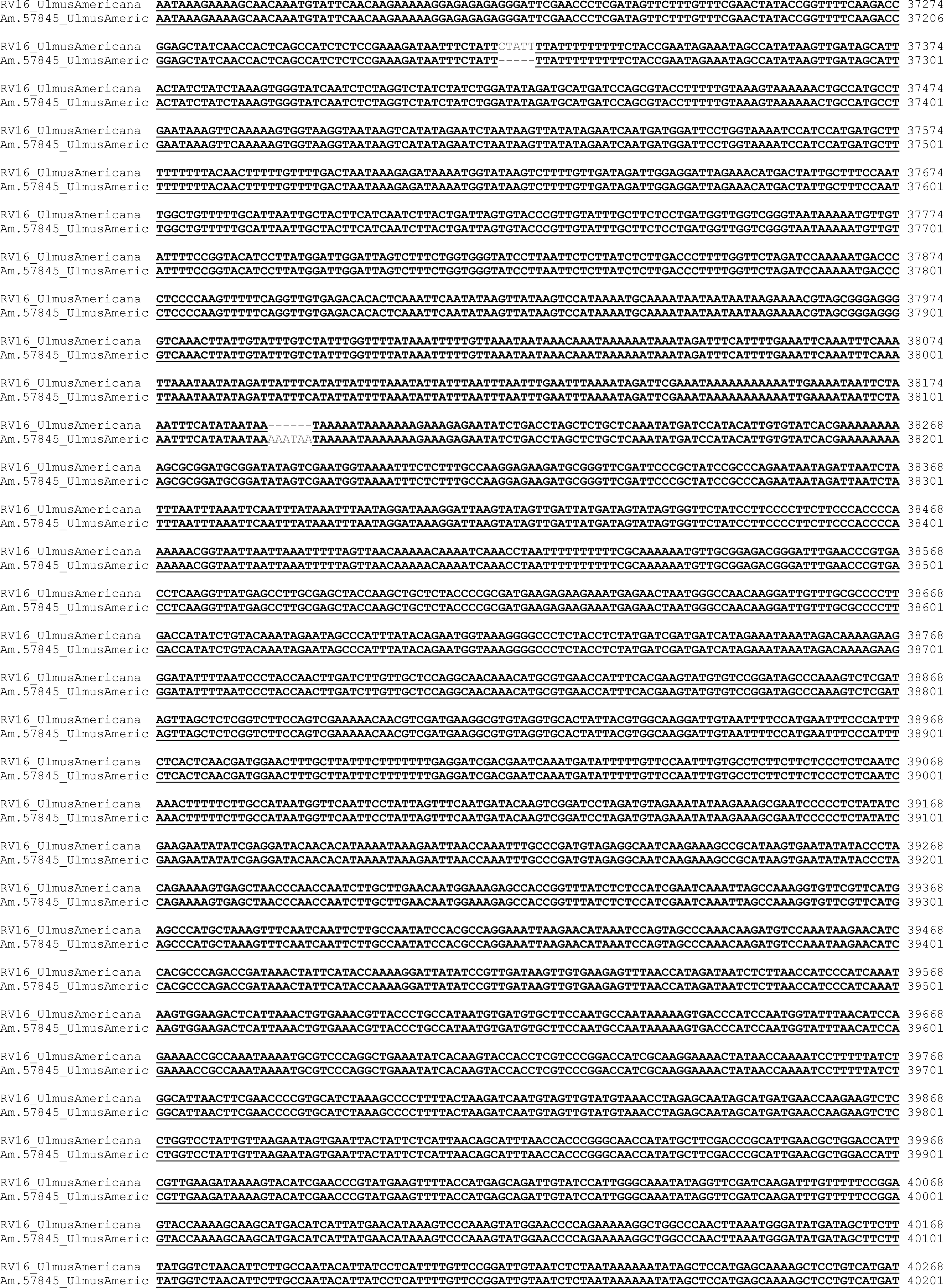
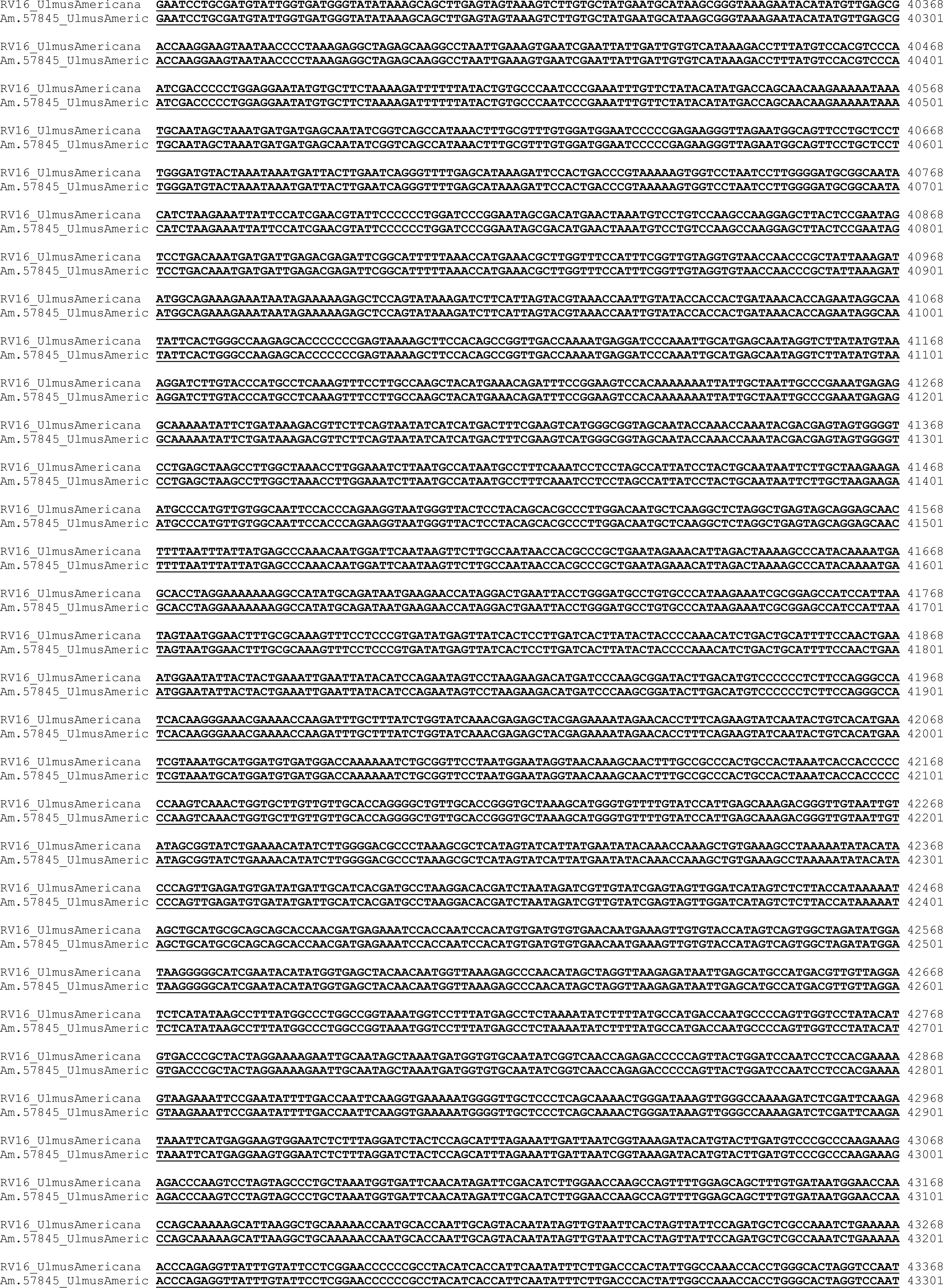
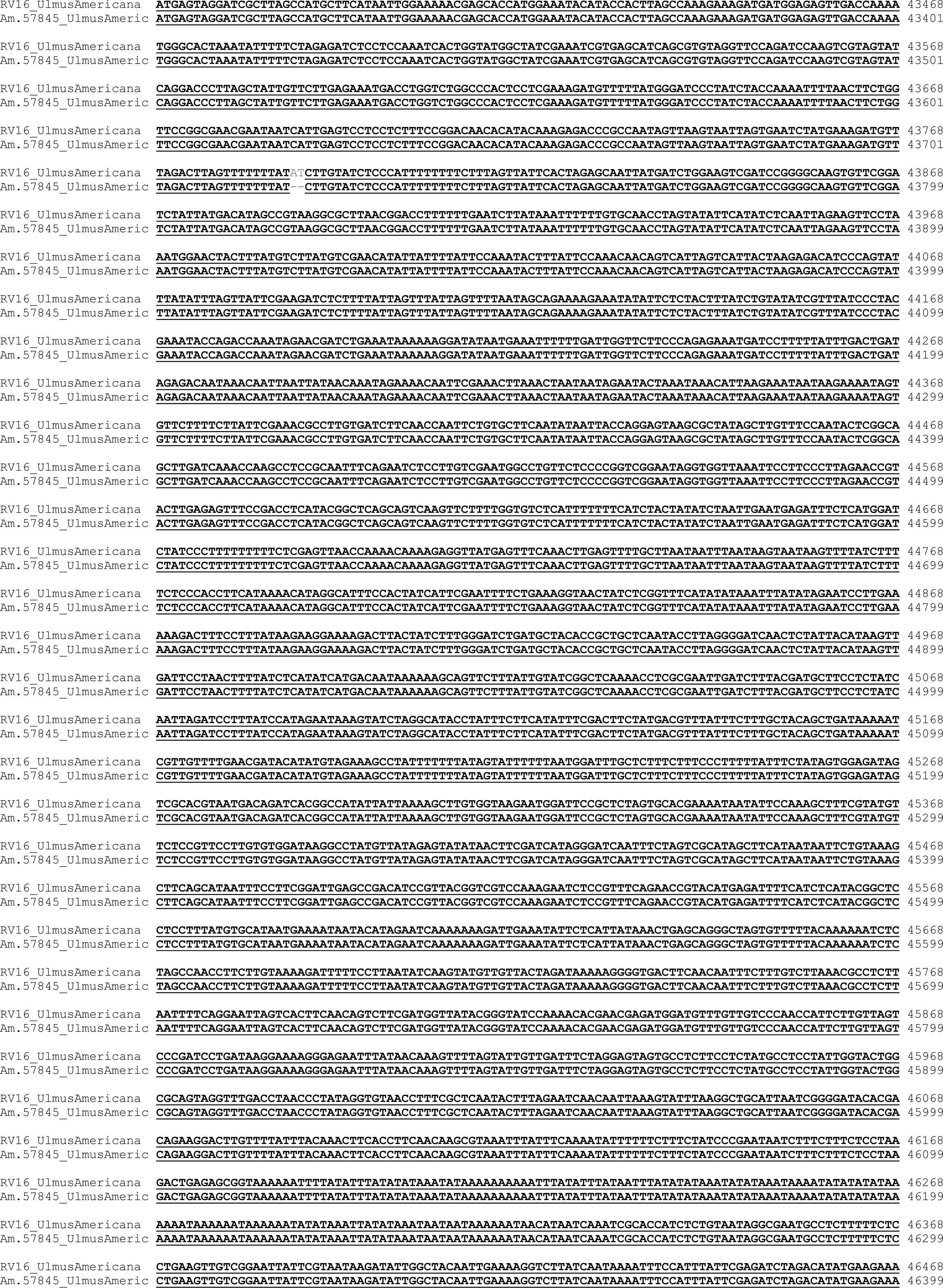
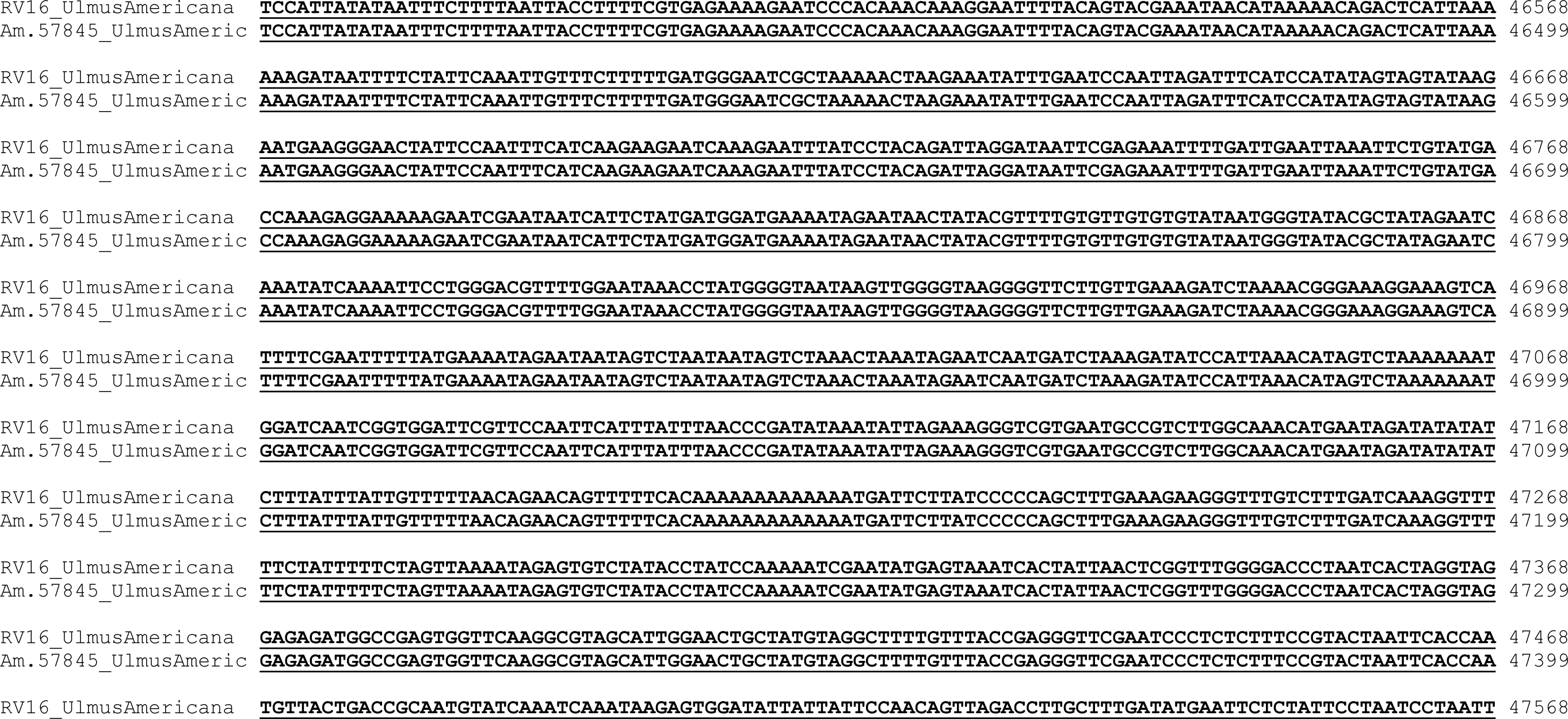

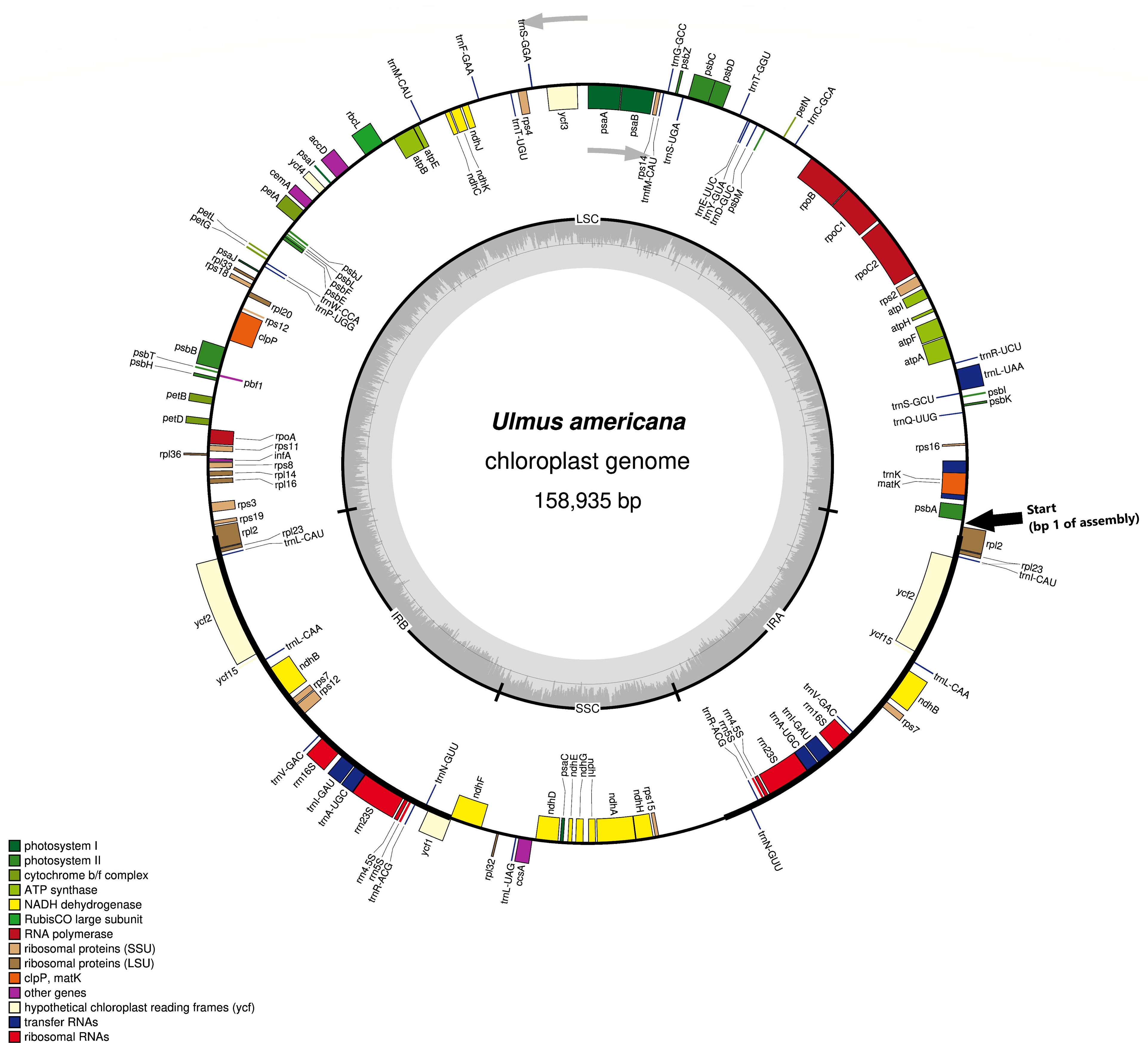

